# Novel Generic Models for Differentiating Stem Cells Reveal Oscillatory Mechanisms

**DOI:** 10.1101/2021.05.28.446119

**Authors:** Saeed Farjami, Karen Camargo Sosa, Jonathan H.P. Dawes, Robert N. Kelsh, Andrea Rocco

## Abstract

Understanding cell fate selection remains a central challenge in developmental biology. We present a class of simple yet biologically-motivated mathematical models for cell differentiation that generically generate oscillations and hence suggest alternatives to the standard framework based on Waddington’s epigenetic landscape. The models allow us to suggest two generic dynamical scenarios that describe the differentiation process. In the first scenario gradual variation of a single control parameter is responsible for both entering and exiting the oscillatory regime. In the second scenario two control parameters vary: one responsible for entering, and the other for exiting the oscillatory regime. We analyse the standard repressilator and four variants of it and show the dynamical behaviours associated with each scenario. We present a thorough analysis of the associated bifurcations and argue that gene regulatory networks with these repressilator-like characteristics are promising candidates to describe cell fate selection through an oscillatory process.

## 1 Introduction

In 1940, Waddington proposed representing the complex regulatory dynamics driving the process of cellular differentiation as an ‘epigenetic landscape’ [1] through which a single cell can be thought of as travelling. The differentiating cell encounters successive ‘decision points’ in the landscape morphology that correspond to differentiation events. These decision points emerge from dynamical changes in the underlying cellular gene regulatory network (GRN), and are mathematically described by variations in parameters that influence the GRN dynamics. For example in zebrafish it is known that Wnt signalling plays a fundamental role in specification and commitment of melanocytes [2–4], and the level of Wnt expression may therefore be considered as a parameter, driving the cellular GRN through one or more of the decision points hypothesised by Waddington.

Despite the philosophical attractiveness of the epigenetic landscape metaphor, the details have remained unclear and no completely self-consistent mathematical description has emerged. For example there are debates regarding the types of bifurcations characterising differentiation: saddle-node bifurcations [5,6] have been proposed as being more consistent with the expected topologies of the core GRN responsible for differentiation than pitchfork bifurcations. Experimental support for these mathematical bifurcation phenomena, e.g. in embryonic stem cells (ESCs), is also lacking. For example, when progesterone is washed away from mature Xenopus oocytes, they do not dedifferentiate but remain mature [22], a finding that is more consistent with the saddle-node bifurcation than with Waddington’s landscape.

Of course, the detailed biological understanding of cell differentiation dynamics has grown significantly recently and it is appreciated that qualitatively different mechanisms may be at work in different contexts. For instance, detailed study of the differentiation of ESCs into two cell types that form either embryonic or extraembryonic tissues indicates that this does not occur in a simple deterministic manner [7]. Instead, cells appear to pass transiently through intermediate states, each seemingly primed to differentiate into one of the possible cell fates [8,9]. Hence ESCs might maintain a collection of meta-stable states, and a better framework to understand and interpret experimental data is required [9].

Most fundamentally, the GRN topologies generating such dynamic stem cell states are poorly understood. In the present work we focus on the question of how broader multipotency (i.e. more than two fates) might generically be generated, in a way that allows differentiation to be biased towards locally-favourable outcomes. We note this is a topic of much current interest in systems biology, particularly noting the exploration of connections between GRN topology and oscillatory dynamics [12], generic links with differentiation dynamics [13–15], and e.g. reshaping the epigenetic landscape by rewiring the GRN using cross-repression among transcription factors (TFs) with self-regulation [14].

In this paper, we analyse and compare five minimally-constructed GRNs that admit cyclical dynamics and allow evolution towards fate commitment controlled by an external stimulus. Our GRNs are variants of the standard repressilator, a simple but fundamental circuit made up of three genes repressing each other. Theoretical modelling of this circuit and its subsequent engineering in *E. coli* have shown its capability to reproduce oscillatory behaviours under broad parameter ranges [17].

We are here interested in adopting this model and the variants proposed to mimick entry into the dynamical oscillatory state characterizing multipotency, and the exit from it, towards a fully differentiated cell state.

Therefore, besides exhibiting oscillations, the system must be equipped with a mechanism to exit the oscillatory regime and proceed towards differentiation. Hence the system must exhibit a sequence of at least two bifurcations, accounting for entering and exiting the oscillatory phase.

We identify two generic scenarios for entering and exiting the oscillatory regime, and we test both in our proposed GRN models. In the first scenario (S1), we hypothesise that a single parameter (i.e. one intervening signalling pathway) drives the system both into and out of the oscillatory regime. An example is Wnt signalling, required for the induction of neural crest, but also for specification of both melanocyte and sensory neuron fates through activation of key TFs (Mitf and Neurogenin, respectively) [3,4,18–23], which we envisage as having this two-fold role. First, increasing Wnt signalling promotes neural crest cells to enter an oscillatory multipotent phase in which fate specification TFs are cyclically expressed; secondly, at higher signalling levels, it promotes the cell’s exit from the oscillatory phase allowing specification of single cell fates through increased Mitf or Neurogenin expression which drive melanocyte or sensory neuron differentiation respectively.

In the second scenario (S2) we hypothesise again that a first bifurcation to oscillatory behaviour is driven by an external signalling pathway, but its influence is then blocked above a critical value. Exit from the oscillatory regime results from other signalling pathways, associated with other parameters. This would arise for a multipotent progenitor that in response to a specific signal starts oscillating, as in S1, driving expression of multiple fate specification TFs, each characteristic of a different ‘sub-state’. Close to each transcriptional sub-state, the stem cell would express distinct cell signalling receptors; then sufficient activation of these receptors by environmental ligands could provide a mechanism that would move the cell out of the oscillatory phase, thus driving differentiation.

The paper is structured as follows. In Section 2, we consider the simplest GRN exhibiting oscillatory behaviour, the so-called repressilator [17]. We find that the system exhibits a parameter regime where trajectories tend towards an attracting limit cycle featuring a slowdown when passing near three sub-states; this appears as a rather weak effect but is numerically detectable. However, exit from the oscillatory regime leads to a single equilibrium, making this GRN unable to select between alternative differentiated states.

This result motivates Sections 3 and 4 where we extend the standard repressilator to include a second repressive circuit opposing the first; we therefore term this GRN the ‘cross-repressilator’. Within this new GRN we consider whether the two TFs act on each gene as either an ‘OR gate’ or an ‘AND gate’, leading to two variants of this circuit. We find that for the ‘OR gate’, the circuit first transitions to oscillatory dynamics, as the standard repressilator, but then followed by a further transition to a stably differentiated cell state, in which only one of the genes is stably expressed. The ‘AND gate’ is even more interesting, since it allows simultaneous expression of two out of the three genes. In both cases our results appear to hold for a large class of modelling choices for these gene interactions.

In Section 5 we extend the ‘AND gate’ cross-repressilator circuit to four and five genes. We show that co-expression of up to three genes is possible depending on the topology of the GRN. These dynamics are compatible with both S1 and S2, as they provide an exit mechanism dependent on a single or multiple bifurcation parameters into a number of alternative equilibria. Furthermore, we note that in all the models, when a cell is in the oscillatory phase, scoring for gene expression of fate-specific TFs would result (in a snapshot view) in the cell being considered ‘fate specified’, despite the fact that it actually retains full multipotency.

A discussion of the biological implications of our models is presented in Section 6.

## 2 The standard repressilator

We first consider the repressilator circuit [17], whose GRN is shown in Fig. 1. Here each gene represents a master regulator transcription factor (TFs) of the differentiation process for a specific fate.

**Figure 1:**
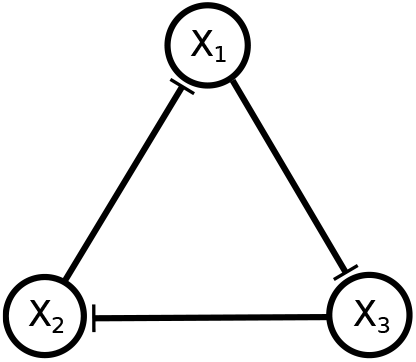
A schematic representation of the repressilator circuit.

Assuming fast (un)binding dynamics of the TFs to DNA, and fast mRNA dynamics, we can reduce the system by adiabatic elimination (see Supplementary Materials) [24], and describe its dynamics with ordinary differential equations (ODEs):

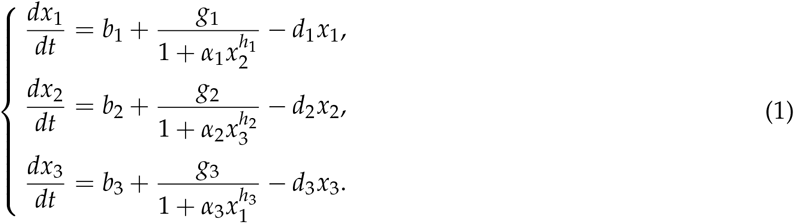

Here, *x_i_* (*i* = 1, 2, 3) describe the concentrations of the proteins X_*i*_, *b_i_* are background expression rates, *g_i_* are maximal gene expression rates (which combine transcription, translation and mRNA degradation rates), *α_i_* are ratios of binding to unbinding rates (association constants) of TFs to DNA, *h_i_* are Hill coefficients describing possible cooperative effects (like multimerisation of the TFs [24]), and *d_i_* are TF degradation rates.

Analysis (see e.g. [17]) shows that the repressilator supports oscillatory behaviour over wide regions of parameter space. Fig. 2 shows (from left) numerical solutions exhibiting the emergence of a family of stable periodic orbits through a Hopf bifurcation as the parameter *g*_1_ increases, time series of sustained oscillations, and the limit cycle in phase space. Oscillatory behaviour exists when *h* > *h*\* ≈ 2.20498, for the parameter values used in Fig. 2.

**Figure 2:**
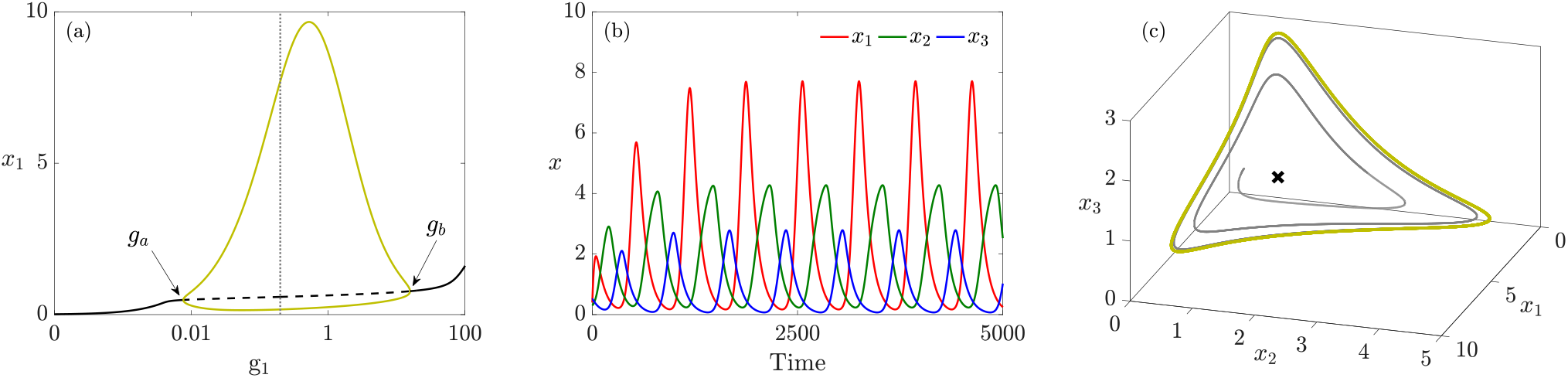
a) Bifurcation diagram for the repressilator as *g*_1_ varies. A family of stable periodic orbits (olive), shown with maxima and minima of *x*_1_, emanates from a supercritical Hopf bifurcation at *g_a_* and terminates at another supercritical Hopf bifurcation at *g_b_*. The stable (solid) branch of equilibria (black) becomes unstable (dashed) between these two bifurcations. The dotted vertical indicates the *g*_1_-value used for panels (b) and (c). b) Time courses of oscillatory concentrations *x*_1_, *x*_2_, *x*_3_ when *g*_1_ = 0.2, *g*_2_ = *g*_3_ = 0.05. c) The solution shown in panel (b) in the phase space (grey) converging to the periodic orbit (olive). The cross (×) is the unique saddle equilibrium, at *x_eq_* = (0.58469, 0.87298, 0.45578). Remaining parameter values are: *b_i_* = 10^−5^, *α_i_* = 50, *d_i_* = 0.01, *h_i_* = 3, for *i* = 1,2,3.

For *h* = 3, the oscillations exist between the two Hopf bifurcations at *g_a_* ≈ 0.00764 and *g_b_* ≈ 15.7919 suggesting that the system is an example of S1 – variations in a single parameter are sufficient to enter and also to exit the oscillatory regime. Our hypothesis for S1 is that external signalling might increase *g*_1_ = *g*_1_(*t*) over time, letting the system sequentially explore non-oscillatory, oscillatory, and a new non-oscillatory regime. However, exit from the oscillatory regime at *g*_1_(*t*) = *g_b_* implies convergence onto the single available equilibrium, making the GRN unable to capture the idea of multiple equilibria necessary to describe stem cell differentiation.

Further, the period of the limit cycles remains bounded as *g*_1_ varies. The shape of the limit cycle (Fig. 2(c)) shows that the amplitudes of the oscillations for different protein concentrations may differ significantly, and generally appear to spike as the orbit approaches the coordinate axes; two proteins are expressed at relatively low levels while the third is higher. Our simulations show that even when the spiking of gene X_1_ is more pronounced, with a huge increase in oscillation amplitude, only a logarithimic or algebraic dependency of the oscillation period appears (see Supplementary Materials), demonstrating the absence of any substantial slow-down around the genes maximal expression substates, and failing thereby to effectively describe any of the scenarios S1 or S2. Furthermore, no mechanism is provided in this model for the stem cell to differentiate into more than one cell type, as no alternative equilibria exist. Thus the standard repressilator does not capture the qualitative changes in behaviour associated with differentiation.

## 3 The cross-repressilator with an ‘OR gate’

We now introduce a variant of the standard repressilator (‘cross-repressilator’), where the inhibitory cycle is accompanied by a second inhibitory cycle arranged in the opposite direction, as depicted in Fig. 3. We assume that regulation of each gene follows an ‘OR gate’, i.e. each gene is active when *both* inhibitors are absent (see Supplementary Materials).

**Figure 3:**
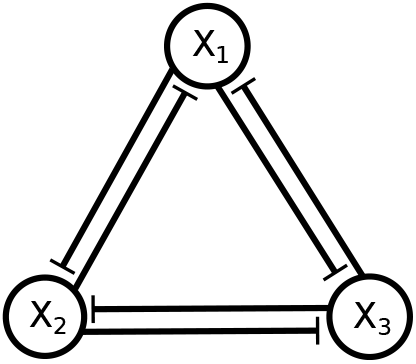
Illustration of cross-repression among three genes X_1_, X_2_ and X_3_ formed with two copies of the circuit shown in Fig. 1, oriented in opposing directions.

Making the same assumptions as the standard repressilator, the dynamics of this GRN can be written as

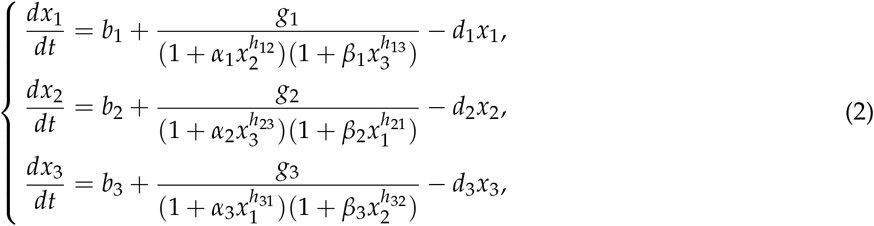

where the repression of each gene is described by the multiplication of two decreasing Hill functions to capture the assumed ‘OR gate’, (see Supplementary Materials). Like system (1), the last term in each equation represents protein degradation. For simplicity, and since it does not cause qualitative changes in the system dynamics, we remove basal expression here and from now throughout this paper. Also, in the numerical simulations we set parameter values equal for each gene, i.e. *g_i_* = *g*, *α_i_* = *α*, *β_i_* = *β*, *d_i_* = *d*, and *h_ij_* = *h* for *i*, *j* = 1,2,3; we return to this in Section 3.2.

Numerical simulations of ODEs (2) are shown in Fig. 4. Figure 4(a) shows the bifurcation diagram for system (2) with respect to the common parameter *g*, all other parameters fixed. The (dashed) solid curves indicate (un)stable equilibria (black) or (maxima and minima of) periodic orbits (olive green). For *g* ⪅ 0.4, system (2) has a single stable equilibrium which is (due to the equalities in parameter values) symmetric: *x*_1_ = *x*_2_ = *x*_3_. The cyclic symmetry apparent in (2) implies that the Jacobian matrix evaluated at this symmetric equilibrium has a pair of eigenvalues with equal real parts (see Supplementary Materials). When this complex conjugate pair crosses the imaginary axis, a supercritical Hopf bifurcation occurs generating a family of stable periodic orbits. As *g* increases further, the stable periodic orbits disappear at a global bifurcation involving three new equilibria created at saddle-node bifurcations.

**Figure 4:**
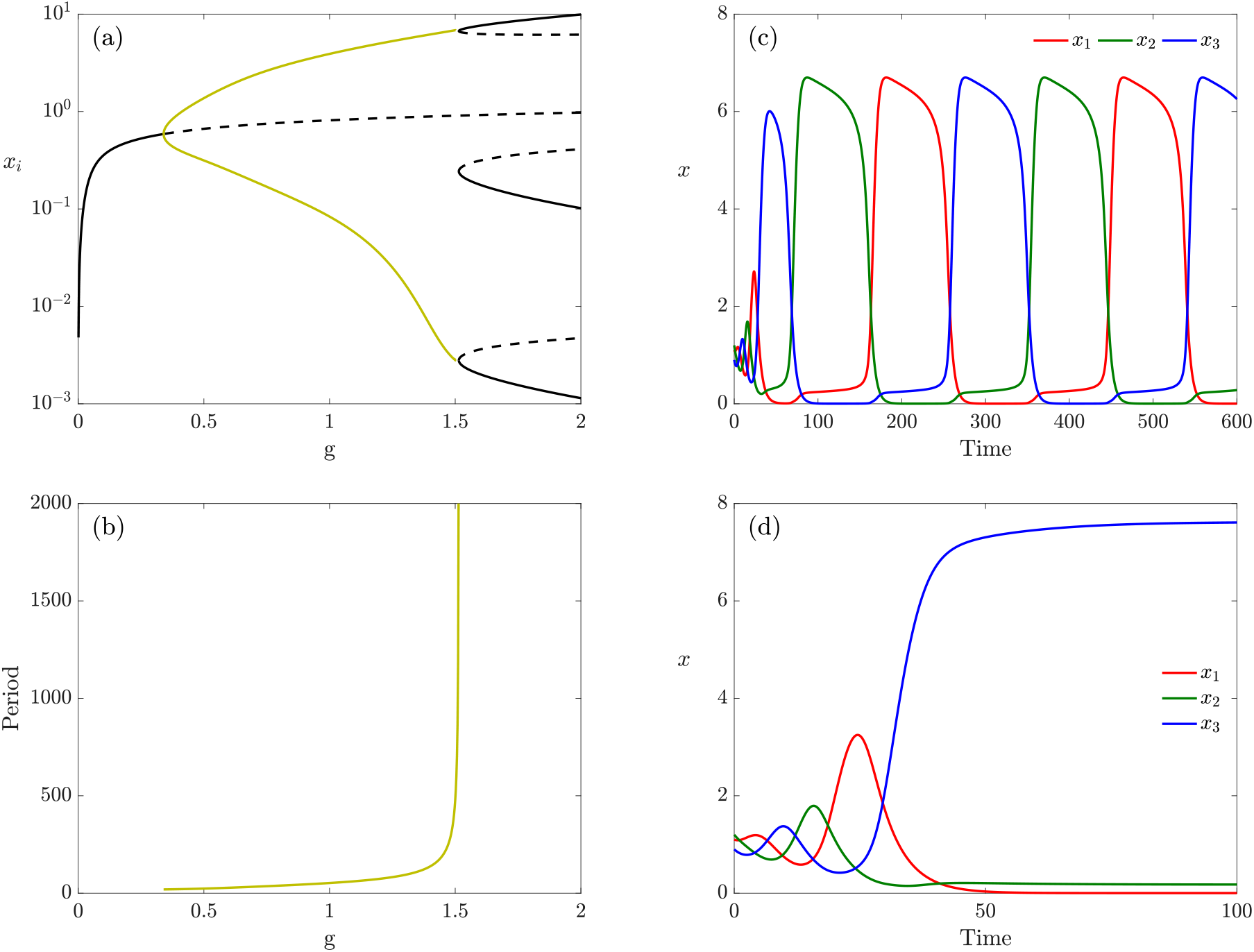
a) Bifurcation diagram with respect to *g* in log-scale for *α* = 9, *β* = 0.1, *h* = 3, and *d* = 0.2. b) The period of the periodic orbits shown in panel (a) increases to infinity while *g* tends to the SNIC bifurcation. c) Time courses of the expression level of genes X_1_ (red), X_2_ (green) and X_3_ (blue) showing the sustained oscillations, for *g* = 1.5. d) For *g* = 1.6, the system stops oscillating and exits to a differentiated cell.

Moreover, there are two distinct global bifurcation mechanisms that remove the periodic orbits. For *α* =9 (shown in Fig. 4), we find that the periodic orbits disappear at a SNIC (Saddle-Node on an Invariant Cycle) bifurcation where three saddle-node bifurcations, related by the cyclic symmetry, occur on the periodic orbit simultaneously. The SNIC bifurcation is well-known in simple dynamical systems, such as the nonlinear pendulum with forcing and damping [25] and in oscillators under external periodic perturbations [26,27]. For *g* >*g*_SNIC_ ≈ 1.51449, system (2) has no periodic orbit but three equilibria related by the cyclic symmetry. For each of the equilibria, one of the *x_i_* is significantly larger than the other two. Initial conditions (ICs) determine which of these equilibria attract the trajectory.

Figure 4(b) shows that, in contrast with the standard repressilator, the period of the periodic orbits diverges as *g* approaches *g*_SNIC_. Typical trajectories spend longer and longer in the vicinity of the three ‘slow regions’ of phase space where the saddle-node bifurcations are about to occur, and exhibit rapid transitions between these ‘slow regions’. This phenomenon has been referred to as the ‘ghost of SNIC’ [30], and indicates the possibility of S2. We identify these slow regions with the sub-states discussed earlier.

In Fig. 4 (right), we show time courses of system (2) before (c) and after (d) the SNIC bifurcation with same ICs. Figure 4(c) shows the time evolution of the three genes for g*g* = 1.5 < *g*_SNIC_. Whenever one of the genes is highly expressed, the expression levels of the other two is low. In Fig. 4(d), we observe the exit from the oscillatory behaviour associated with a further increase of parameter *g* (*g* = 1.6). Here, gene X_1_ is stably expressed at a high level, while genes X_2_ and X_3_ are stably expressed at very low levels.

In summary, for *g* increasing, the system shows three different behaviours: (i) convergence to a (unique) stable equilibrium; (ii) oscillations among three sub-states; and (iii) attraction to one of the stable equilibria. The natural interpretation, compatible with S1, is to see the first two cases as reflecting multipotency within the stem cell, while the third case represents attainment of one of multiple differentiated cell types.

The phenotype attained in the differentiated state depends on the IC used for the simulation and its intrinsic properties, here illustrated through different *g*-values, as shown in Fig. 5. Figure 5(a) colour-codes the (*x*_1_, *x*_2_)-plane according to the final equilibrium for three different ICs for *x*_3_(*t*), and fixed *g*. Figure 5(b) shows the effect of varying parameter *g* on the same colourmap when *x*_3_(0) is fixed. The three coloured regions in the (*x*_1_, *x*_2_)-plane twists around a central point. Increasing *x*_3_(0) tightens the twists of these spiralling regions while the opposite occurs when *g* increases. Moreover, variations in *g* keep the central twisting point fixed at (*x*_1_(0), *x*_2_(0)) = (1,1) whereas increasing *x*_3_(0) makes this point drift away.

**Figure 5:**
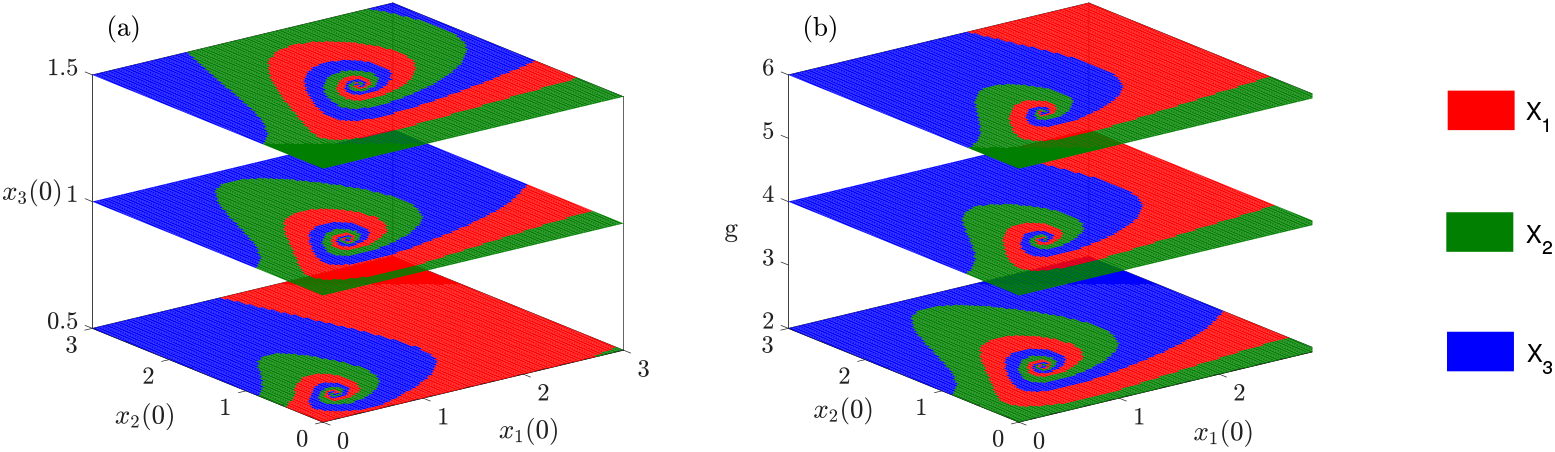
a) Colourmaps indicating which gene is expressed starting from ICs in the (*x*_1_, *x*_2_)-plane, for *g* = 2 and three different ICs of *x*_3_: *x*_3_(0) = 0.5,1 and 1.5. b) The same colourmaps as panel (a) for *x*_3_(0) = 1 and three different values of *g*: *g* = 2, 4 and 6.

To obtain insight into the feasibility of S2 for this system, we explore the bifurcation behaviour in the (*g*, *α*)-plane. Fig. 6(a) shows the continuation of the bifurcations identified for *α* = 9 in the (*g*, *α*)-plane. When *α* decreases, the location of the Hopf bifurcation curve (HB) moves to larger *g* whereas the curve of saddle-node bifurcations (SN) on which the SNIC bifurcation occurs moves to lower values of *g*. At *α* = *α*\* ≈ 3, the SNIC regime terminates at a codimension-two bifurcation, indicated by a grey dot. Two new bifurcation curves emerge from this point, indicated by the labels HC (homoclinic) and SN (saddle-node). For *α* < *α*\* the three saddle-node bifurcations (due to symmetry) occur away from the periodic orbit in phase space, resulting in a region of the (*g*, *α*)-plane in which these new equilibria (three stable and three saddle equilibria) coexist with the stable periodic orbits. The periodic orbits then collide with the three saddle equilibria at a global bifurcation and disappear. As in the SNIC bifurcation, the period of the periodic orbits goes to infinity as the orbits approach the global bifurcation, but the mathematical details of the behaviour differ from the SNIC case, as shown for example by the scaling of the period with distance to the bifurcation point being different (see Supplementary Materials).

**Figure 6:**
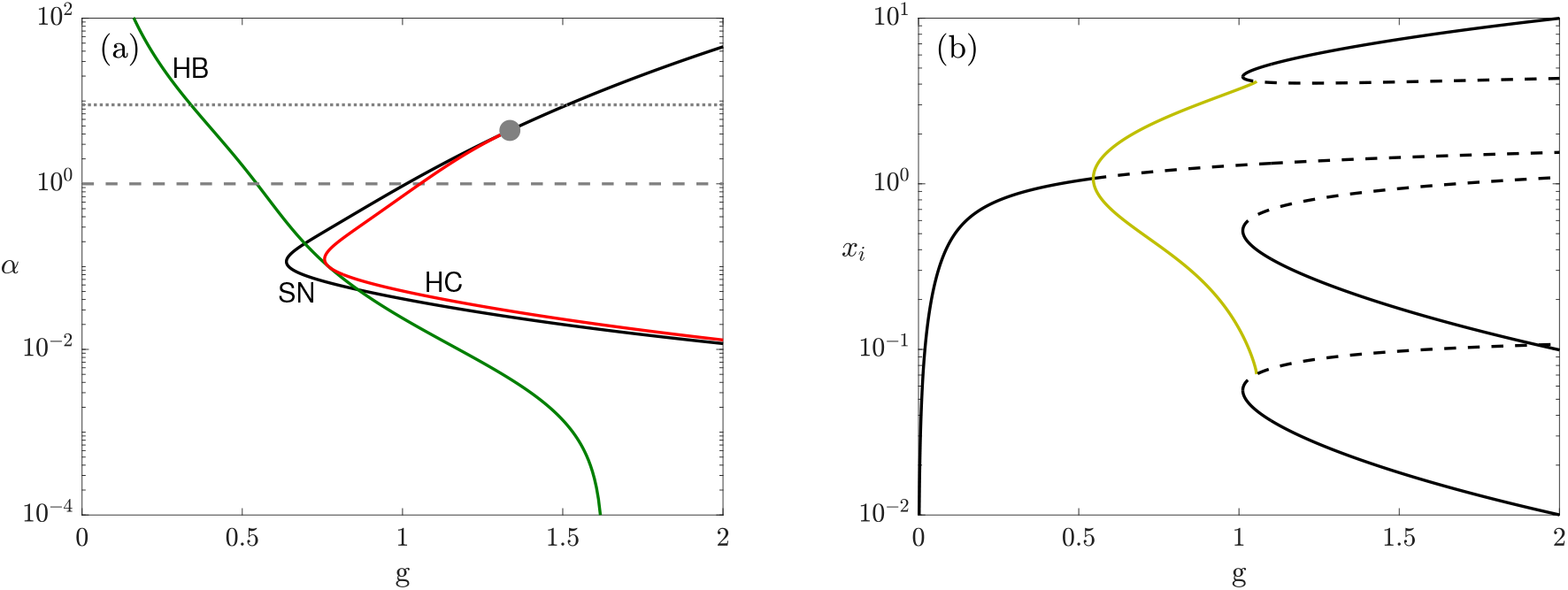
a) Two-parameter continuation of saddle-node (black), Hopf (dark green) and heteroclinic (red) bifurcations, shown in Fig. 4(a), in the (*g*, *α*)-plane; vertical axis (*α*) is shown in log-scale. The dotted and dashed lines indicate the corresponding *α* values used in Fig. 4(a) (*α* = 9) and in panel (b) (*α* = 1), respectively. b) Bifurcation diagram of system (2) with respect to g in log-scale when *α* = 1. The diagram structure is mainly similar to the one in Fig. 4(a). However, here, a small region of multi-stability is created between the oscillatory regime and the stable equilibria such that the family of periodic orbits terminates at a heteroclinic bifurcation (instead of a SNIC).

We observe also that as *α* decreases further the heteroclinic bifurcation (HC) and SN curves move further apart from each other in the (*g*, *α*)-plane, with maximum separation at *α* ≈ 0.1.

For *α* < *α*\* the HB (green) and HC curves (red) determine the left and right boundaries of the oscillatory region, respectively. Also note that the HB and HC curves meet and are tangent to each other at *α* = 0.1 which is not generic but arises here due to the additional symmetry existing when *α* = 0.1 = *β*. When *α* = *β*, ODEs (2) are symmetric under two independent symmetry operations: the cyclic permutation symmetry (*x*_1_, *x*_2_, *x*_3_) → (*x*_2_, *x*_3_, *x*_1_) and the interchange of any two of the three variables, e.g. (*x*_1_, *x*_2_,*x*_3_) → (*x*_2_,*x*_1_,*x*_3_). In group theory notation, the symmetry group is now the symmetries of an equilateral triangle *D*_3_ rather than just the cyclic group of order three, 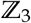. This additional symmetry implies the tangency between the two curves. The analysis around this point will be discussed in detail in a separate paper [32] and has been observed in other contexts [28,29].

Figure 6(b) shows the bifurcation diagram, varying *g*, for *α* = 1 corresponding to the lower horizontal dashed line indicated in Fig. 6(a). Here we observe an interval of *g*-values in which the stable periodic orbit coexists with the three stable equilibria. Consequently we would expect that this transition to a stable equilibrium occurs rather smoothly since the sub-states already exist prior to the termination of the oscillations.

These observations support S2. Once the system has entered the oscillatory regime by tuning of *g* through a first signalling pathway, a second signalling pathway might intervene to lower *α*, stop the oscillations, and drive the system to differentiation. This scenario requires that the change of *α* happens faster than the permanence of the system close to a selected sub-state. This slowdown of the cycling dynamics is therefore essential for this mechanism.

Thus, after establishment of the stem cell, increasing the first signal initiates the cyclical expression of fate-specific TFs corresponding to the different differentiated cell-types. Subsequently, in each primed sub-state, as a result of TF specific activation of sensitivity to fate specification signals, the cell is sensitised to local signals. When a cell receives such a signal for sufficient time, this shifts *α* and removes the oscillations, thereby initiating differentiation.

### 3.1 Illustrating the two scenarios with time-dependent parameters

So far we have assumed that all the parameters of system (2) are fixed and do not vary in time. In this section, we explore more directly the two scenarios S1 and S2 by assuming that relevant parameters are regulated in time by an external signalling pathway. We will consider the case when either one or both of *g* and *α* are time dependent. In S1, parameter g only is responsible for shifting the cell from the multipotent regime into the oscillatory regime and then into the differentiated cell. Accordingly, we assume that *g*(*t*) follows the form of an increasing Hill function, from 0 and saturating at *ĝ* > *g*_SNIC_ as

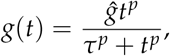

where we set *ĝ* = 1.7, *p* = 1.25 and *τ* = 1000. The bottom panel of Fig. 7(a) illustrates the time-variation of *g*(*t*). The top panel in Fig. 7(a) shows how the system responds to *g*(*t*), moving through the three different regimes as *g*(*t*) increases and tends to the value *ĝ* (compare with Fig. 4(a)). In detail, the system first settles quickly from the IC to the single stable equilibrium and follows it as *g*(*t*) slowly approaches the Hopf bifurcation. After Hopf we observe the onset of rapid but finite-frequency oscillations. As *g* → *g*_SNIC_, the period of oscillations increases significantly, giving rise to dynamical phases close to each of the three sub-states. Finally, when *g*(*t*) crosses *g*_SNIC_, the oscillations disappear and the system settles at one of the three equilibria, each corresponding to a different expressed gene. For example, in Fig. 7, gene X_3_(blue) is expressed and the other two genes remain at a low level.

**Figure 7:**
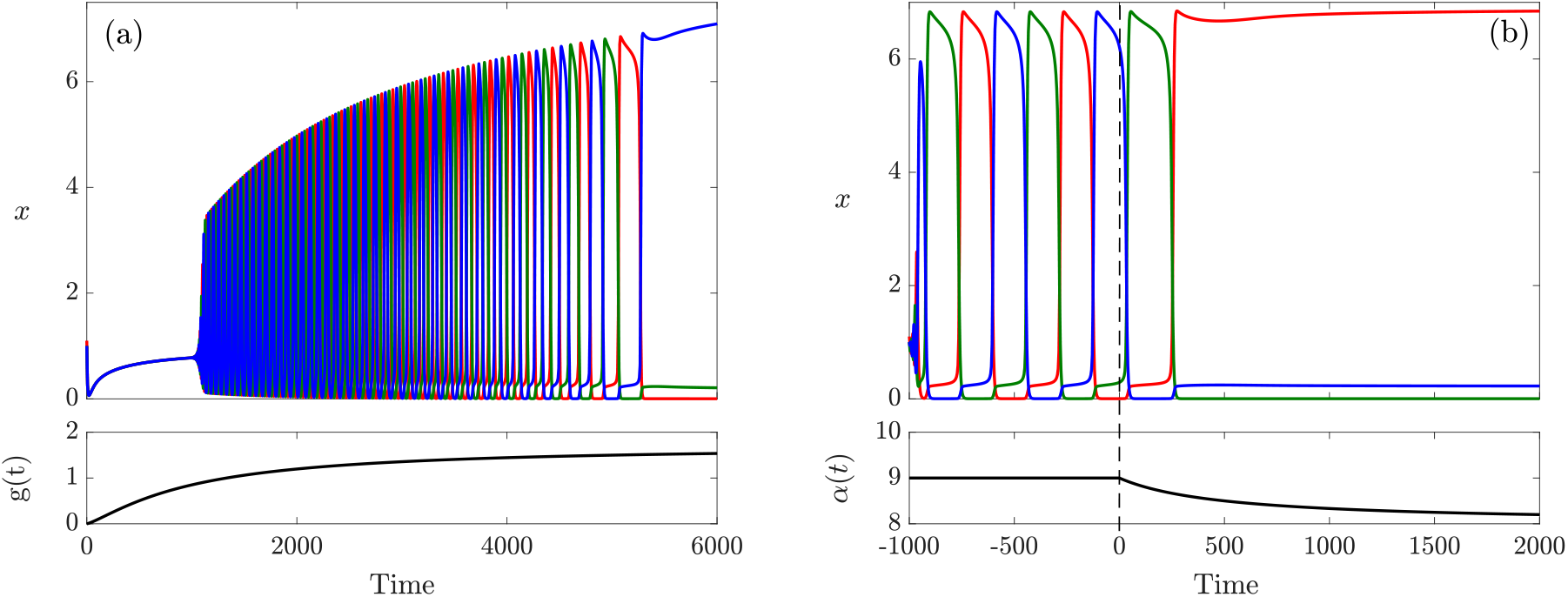
a) The development over time from a multipotent cell to a differentiated cell controlled by a time-dependent profile *g*(*t*) (shown in the lower panel) according to S1. Parameters are as Fig. 4(c). b) Development from a stable oscillatory regime into a differentiated cell according to S2, when the secondary parameter *α* starts decreasing at 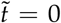 (shown in the lower panel). Here *g* = 1.5 is held constant, and all other parameters take the same values as in Fig. 4(c).

As discussed for S1, the precise state of the system when *g* goes through *g*_SNIC_, as well as the value of the saturation level *ĝ* play crucial roles in selecting cell fate. The shape of the profile of *g*(*t*) is also relevant for this selection.

For S2, we hypothesise that two parameters, *g* and *α*, are involved in cell differentiation. As shown in Fig. 7(b), we first assume that parameter *g* shifts the system into the oscillatory regime while *α* remains fixed. At a certain time, we suppose a second pathway causes a decrease in *α*, modelled as

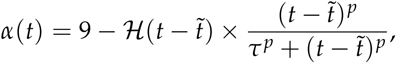

where 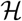 is a Heaviside step function (i.e. 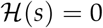 when *s* < 0 and 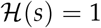 when *s* >0), 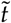 is the time when the time dependency of *α*(*t*) starts, and the constants *τ* = 1000 and *p* = 1 control the profile shape.

S2 clearly allows more control over the final fate of the cell through the increased number of coefficients involved in specifying the time-dependency of the second bifurcation parameter. To understand these further dependencies, we analyse the effect of variations in the switch-on time 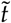 and the time scale parameter *τ* for the saturation of *α*(*t*). Shown in Fig. 8, we colour-code the parameter plane (*τ*,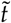) based on the expressed gene for *g* = 1.5 (a) and *g* = 1.51 (b). The (*τ*,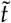)-plane is divided into stripes which periodically alternate between red, green and blue. The lower value of *g* in Fig. 8(a) corresponds to narrower stripes as a result of the smaller oscillation period, with the system being more sensitive to changes in switch-on time and saturation time scale. Further, more horizontal stripes in Fig. 8(b) indicate that cell fates become less sensitive to *τ* for this larger value of *g*, at least in the regime close to the global bifurcation.

**Figure 8:**
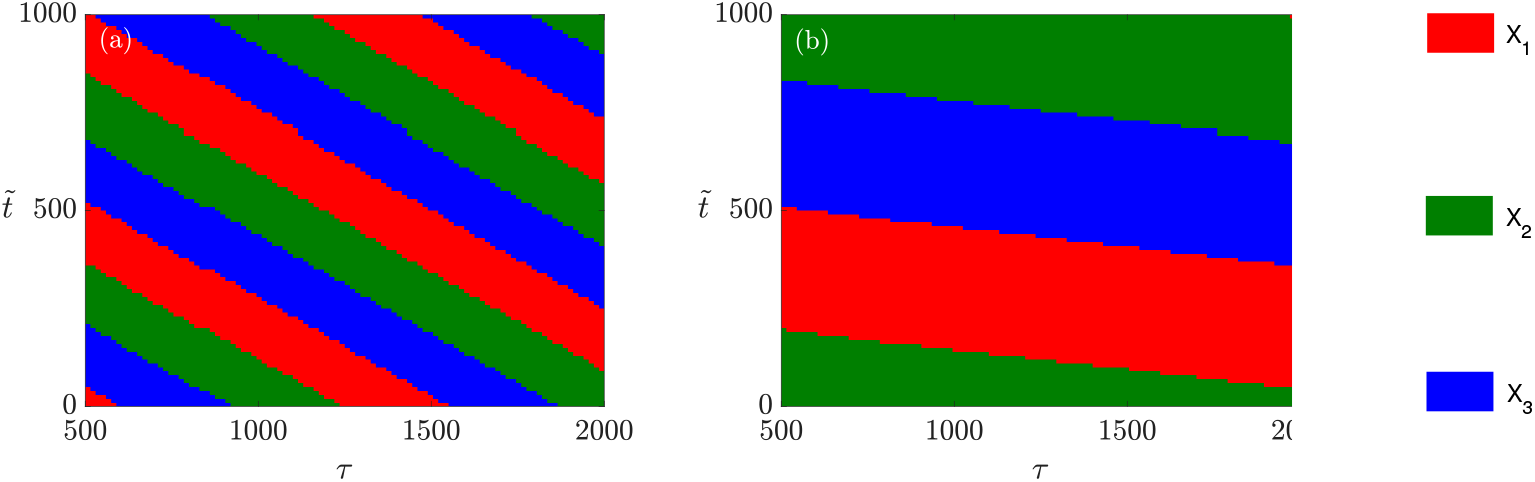
Colourmaps show the expressed gene in (*τ*,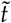)-plane starting from the same IC for *g* = 1.5 (a) and *g* = 1.51 (b).

### 3.2 Breaking the symmetry

Until now, we assumed that system (2) is symmetric and all the association rates *α_i_* and *β_j_*, *i, j* = 1,2,3 for the TFs X_1_, X_2_ and X_3_ are equal. Here, we break the symmetry in the reciprocal repressing loops in Fig. 3 and consider the case that *α_i_* and *β_j_* take different values, as specified in Fig. 9. Note that we keep the remaining parameters *g* and *d*, unchanged.

**Figure 9:**
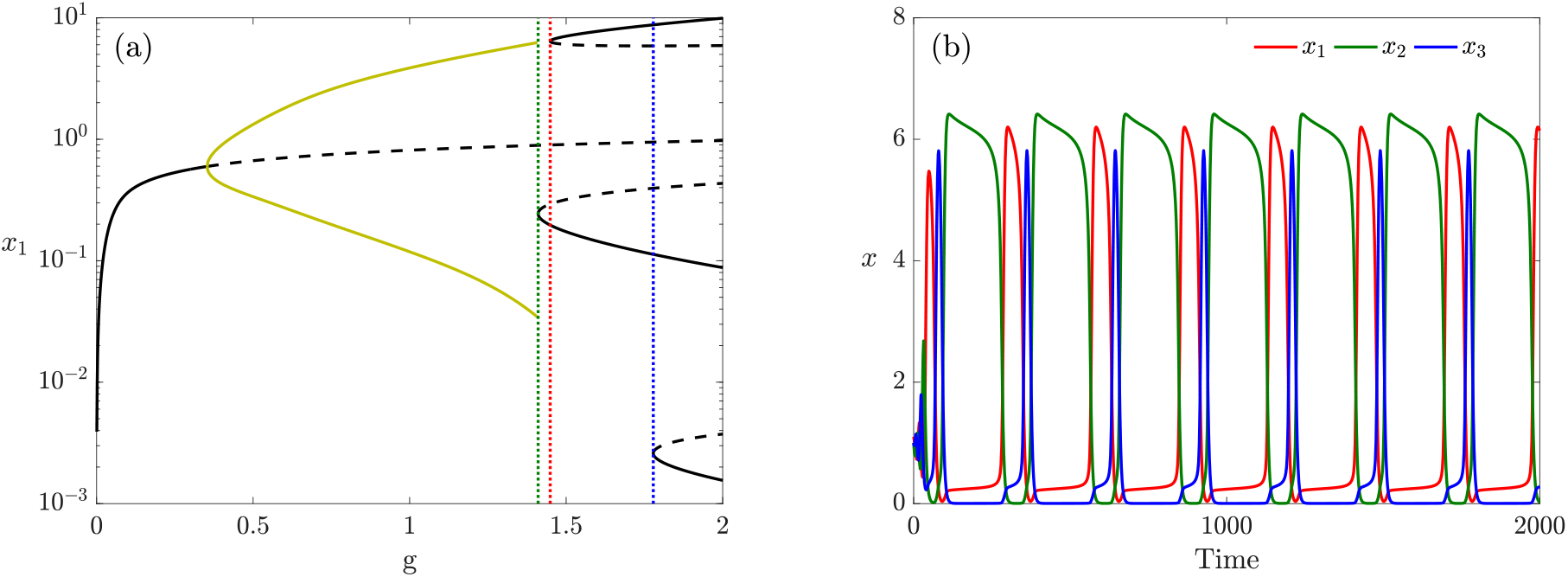
a) Bifurcation diagram of the system for *α*_1_ = 7, *α*_2_ = 9, *α*_3_ = 8, *β*_1_ = 0.115, *β*_2_ = 0.07, *β*_3_ = 0.1 and *d* = 0.2 in log-scale. b) Time courses of the oscillations in the expression level of genes X_1_ (red), X_2_ (green) and X_3_ (blue) for the same parameters in panel (a) and *g* = 1.4.

Figure 9(a) shows the bifurcation diagram with respect to *g* in log-scale. The structure remains qualitatively unchanged compared to the symmetric case. For *g* small, a unique stable equilibrium exists, which undergoes a Hopf bifurcation and becomes unstable. For *g* large, there are six branches of equilibria; the stable branches merge with the saddle-type ones and disappear as *g* decreases. The family of stable periodic orbits emerging from the Hopf bifurcation terminates at a SNIC bifurcation. However, here, only one equilibrium lies on the invariant cycle contrary to the symmetric case, with three equilibria on the invariant cycle. This happens because the saddle-node bifurcations, at which the six branches of equilibria appear, occur at different *g*-values (they are no longer symmetry-related), indicated by the vertical dotted lines.

Figure 9(b) shows time courses of the genes concentrations just before the SNIC bifurcation at *g*_SNIC_ ≈ 1.41054; the other parameter values are as in Fig. 9(a). As before, each TF is expressed cyclically when the other two are at low levels. However, gene X_2_ remains expressed for a significantly longer time than the other two, particularly X_3_. Due to the occurrence of the saddle-node bifurcation with high level of X_2_ at smaller values of *g*, the slow region created around the bifurcation has a stronger effect which leads to a longer ‘dwelling time’ at expressed X_2_. In fact, the dwelling time for each expressed gene depends on the distance of g-value at which the corresponding saddle-node bifurcation (indicated with a dotted vertical line of the same colour) occurs.

Analysis of Fig. 9 determines that with the current parameters, the conditions favour gene X_2_; therefore, when *g* crosses to the right of the SNIC bifurcation, the system settles at a stable equilibrium where X_2_ is expressed. Changing this condition results in a shift in the location of the saddle-node bifurcations along the horizontal axis and might alter the order of saddle-node bifurcations; consequently, lengthening the dwelling time for another gene.

## 4 The cross-repressilator with an ‘AND gate’

The third model we consider here is similar to system (2) with the difference that each gene is repressed only if all other genes are highly expressed (a so-called ‘AND gate’).

This model is described by

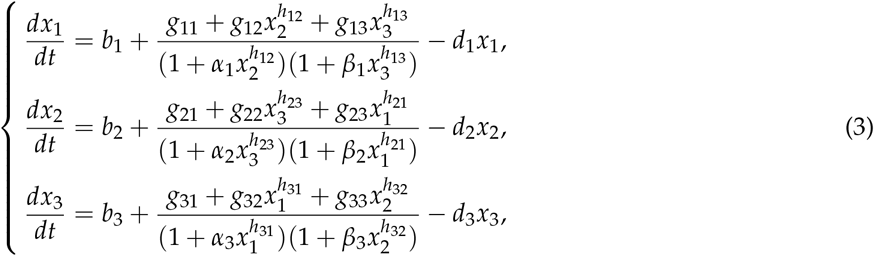

where again the decreasing Hill functions describe the effect of repressing genes (see Supplementary Materials). We also set *g_ij_* = *g_j_* for *i, j* = 1,…, 3 and *d*_1_ = *d*_2_ = *d*_3_ = *d*. We also fix *α_i_* = *β_i_* = 1 and the Hill function exponents *h_i,j_* = 3 for *i, j* = 1,2,3.

As previously, we explore system (3) numerically and compute the bifurcation diagram, varying *g*_1_ while keeping *g*_2_ and *g*_3_ fixed (and unequal). Figure 10 (a) shows a bifurcation diagram for system (3) with respect to *g*_1_. For small values of *g*_1_, the system has a single stable equilibrium (black solid line) which undergoes a supercritical Hopf bifurcation and becomes unstable (dashed line). The stable periodic orbits (olive) are indicated by the maximum and minimum values of *x*_1_. The periodic orbit, similar to the ‘OR gate’ case, terminates at a SNIC bifurcation where it collides with a saddle-node bifurcation. Note that these branches are part of a single branch of equilibria terminating at another oscillatory regime at very large parameter values *g*_1_ (not shown). Panels (b) and (c) in Fig. 10 show time courses of the expression level of each gene in system (3) for values of *g*_1_ mentioned in the caption.

**Figure 10: :**
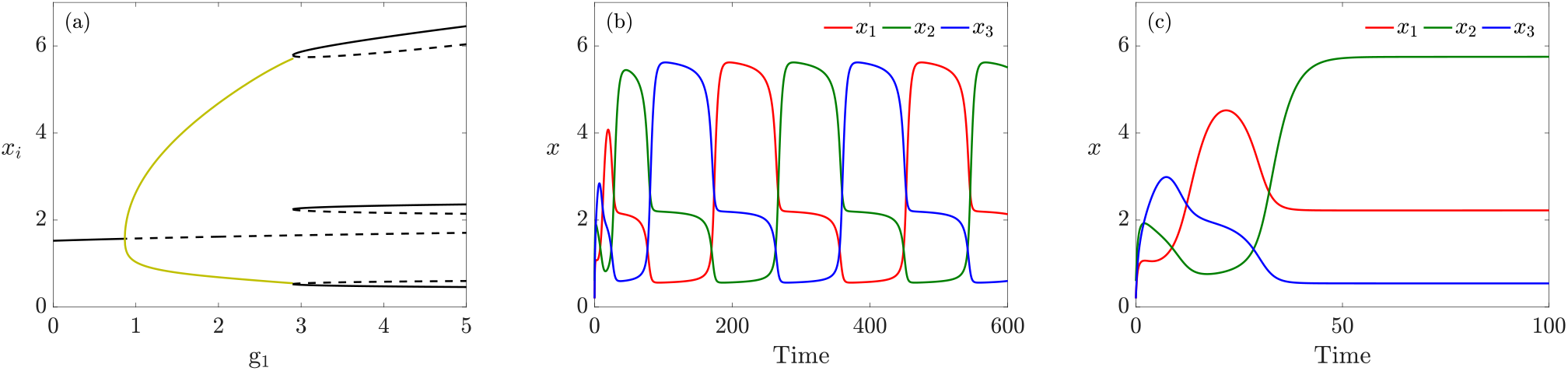
a) Bifurcation diagram of system (4) with respect to *g*_1_ for *g*_2_ = 0.76, *g*_3_ = 1.9 and *d* = 0.3. b) Time courses of the expression level of genes X_1_ (red), X_2_ (green) and X_3_ (blue) showing the emergence of an intermediate expression level in the oscillatory regime of system (3) when *g*_1_ = 2.8. c) Gene expression levels converging to equilibrium in the tristable regime for *g*_1_ = 3.

Clearly, the behaviour of the ‘AND gate’ model follows closely the analysis presented for system (2), but with one important and biologically relevant difference. Compared to system (2), in which only one gene could be highly expressed and the other two genes were necessarily at very low levels, in system (3), two genes can be expressed at high or moderate levels simultaneously. Of course, the precise levels depend on parameter values, but the ‘AND gate’ structure is crucial in admitting states with more than one high expression level. The form of the periodic orbit also changes and shows that it lingers in sub-states where two gene expression levels are nonzero. This notable feature is potentially useful in matching models with quantitative experimental data where it allows one to select a model that allows or prevents the existence of phenotypes dependent upon the expression of a single or a combination of master regulator TFs.

## 5 Extension to four and five genes

So far, we considered GRNs with only three genes. We now investigate the dynamics with more than three genes. For the same cross-repressilator in ‘AND gate’ configuration discussed in the previous section, an odd number of genes is necessary to create a negative feedback loop that supports sustained oscillations [31]. Hence a trivial extension of the cross-repressilator to four genes, preserving the cyclical topology, fails to reproduce oscillatory dynamics, and therefore we do not consider it here. Even numbers of genes in the cycle result in states where the gene expression levels alternate around the loop. However, adding cross connections to this four-gene network so as to build three-gene negative loops, embedded within the four-gene GRN as shown in Fig. 11(a), recovers the oscillatory dynamics. For five genes, the simple cyclical topology of the cross-repressilator, with only ‘nearest neighbour’ interactions and without additional cross-regulation, shown in Fig. 11(b), maintains oscillatory behaviour.

**Figure 11:**
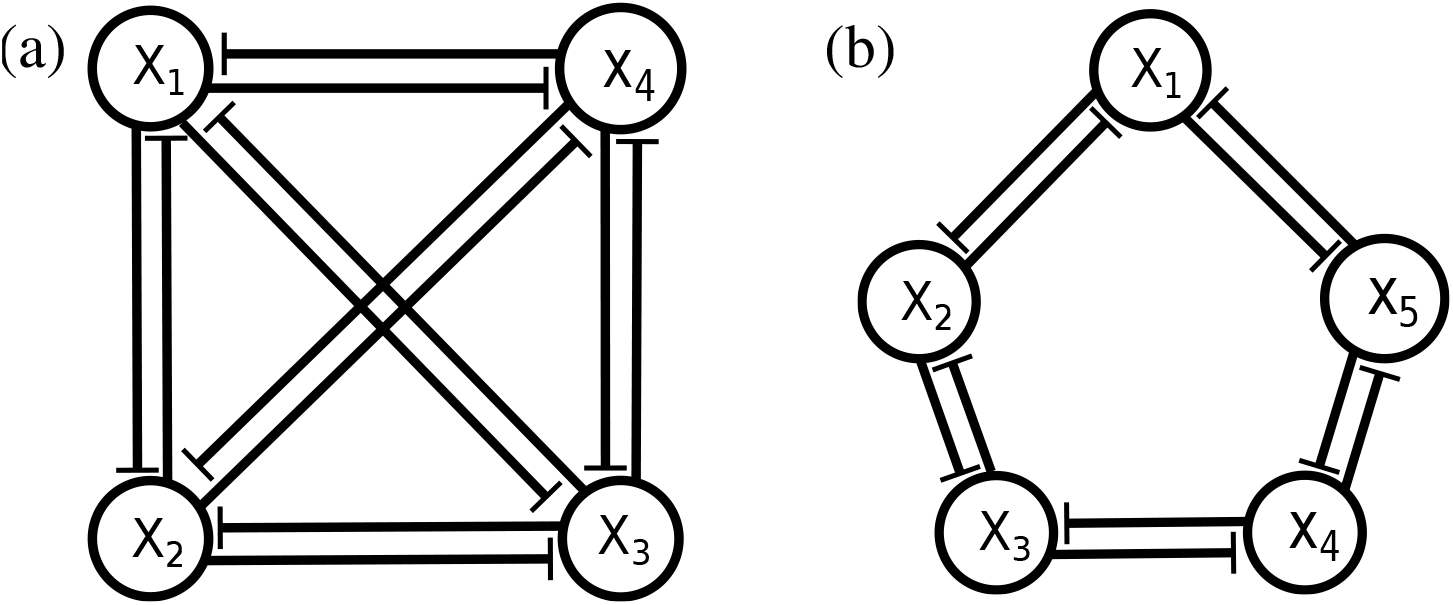
a) Extension of the cross-repressilator GRN to four genes with cross connections that support oscillations. b) A cyclical arrangement of five genes, sufficient to drive oscillations without additional cross connections.

### 5.1 The four gene network

Here, we describe the dynamics of a GRN with four genes, as illustrated in Fig. 11(a). The network is fully connected such that each gene is repressed by the other three genes, operating in ‘AND gate’ configuration. The results in (see Supplementary Materials for details):

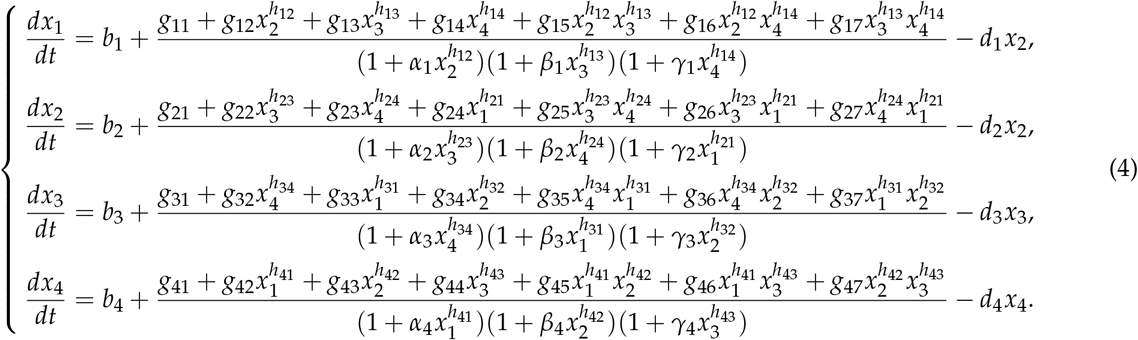

For simplicity, we again set *g_ij_* = *g_j_* and *d_i_* = *d* for *i, j* = 1,…,4. We also keep fixed the values *g_j_* = 0.5 for *j* = 5,…,7, *α_i_* = *β_i_* = *γ_i_* = 1 and *h_ij_* = 3 for *i, j* = 1,…,4 (*i* ≠ *j*).

Figure 12(a) shows the bifurcation diagram of system (4) with respect to *g*_1_, together with time courses of the expression levels of all genes in the system for two specific values of parameter *g*_1_ in the multipotent (b) and differentiated (c) regimes. In Fig. 12(a), a single branch of stable equilibria (solid; black) undergoes a Hopf bifurcation and becomes unstable/saddle (dashed). The envelope of stable periodic orbits (olive) emanating from the Hopf bifurcation terminates at a SNIC bifurcation where new branches of equilibria appear. Four of these new branches are stable, and collide with the other four unstable branches, disappearing at saddle-node bifurcations.

**Figure 12:**
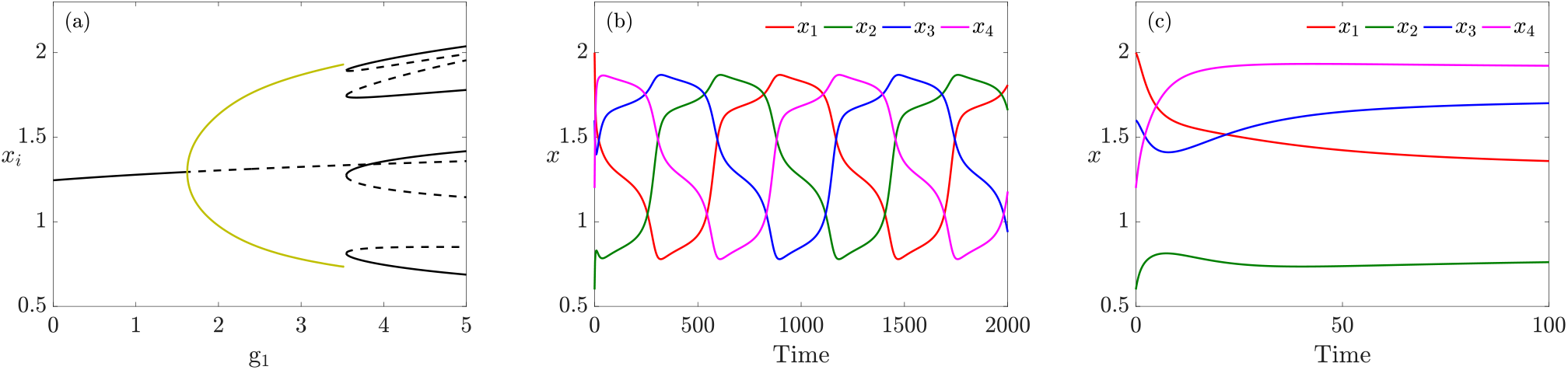
a) Bifurcation diagram of system (4) with respect to *g*_1_ for *g*_2_ = 1.6, *g*_3_ = 0.5, *g*_4_ = 1.5 and *d* = 0.4. (b) and (c) Time courses of the expression level of genes X_1_ (red), X_2_ (green), X_3_ (blue) and X_4_ (magenta) with two different behaviours: in the multipotent oscillatory regime of system (4) when *g*_1_ = 3 (b) and in the multistable regime for *g*1 = 3.6 (c).

Panels (b) and (c) in Fig. 12 show time series of the expression levels of the genes in the oscillatory and differentiated regimes for *g*_1_ = 3 and *g*_1_ = 3.6, respectively. One notable, and biologically relevant, feature of this model is that all four genes can be expressed, with one gene highly expressed, two genes expressed at intermediate levels, and one gene only slightly expressed. This scenario is consistent with differentiation reflecting a combinatorial role for multiple TFs. The relative concentrations of the expressed genes would open up possibilities for differentiation to multiple cell types.

### 5.2 The five gene network

We now consider the five-gene network depicted in Fig. 11(b). The network comprises two inhibitory loops arranged in opposing directions so that gene *X_i_* is repressed by genes *X*_*i*−1_ and *X*_*i*+1_, again in ‘AND gate’ configuration. Therefore, similar to the three-gene case, the dynamical equations are

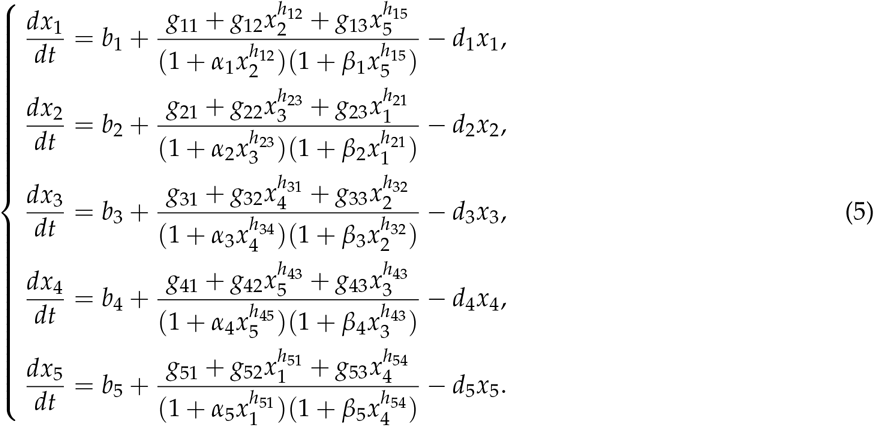

where the maximal expression rates imposed by the anticlockwise and the clockwise loops respectively equal to *g*_1_, *g*_2_ and *d_i_* to *d*. We also fix the values *α_i_* = *β_i_* = *γ_i_* = 1 and *h_ij_* = 3 for *i, j* = 1,…, 5 (*i* ≠ *j*). Figure 13(a) shows the bifurcation diagram remains unchanged with respect to *g*1 except for the number of stable (solid; black) and saddle (dashed; black) branches in the multistable regime. Figure 13(b) exhibits time courses of gene expression levels when *g*_1_ = 2. Figure 13(c) shows gene concentrations for *g*_1_ = 2.5 when the cell differentiates. In the differentiated regime, the number of expressed genes increases to three compared with the three-gene case. We conjecture that more generally for a cyclic network of *n* genes, 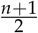 genes are expressed at high or moderate (but unequal) levels and 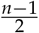 are either not expressed at all or are only weakly expressed.

**Figure 13:**
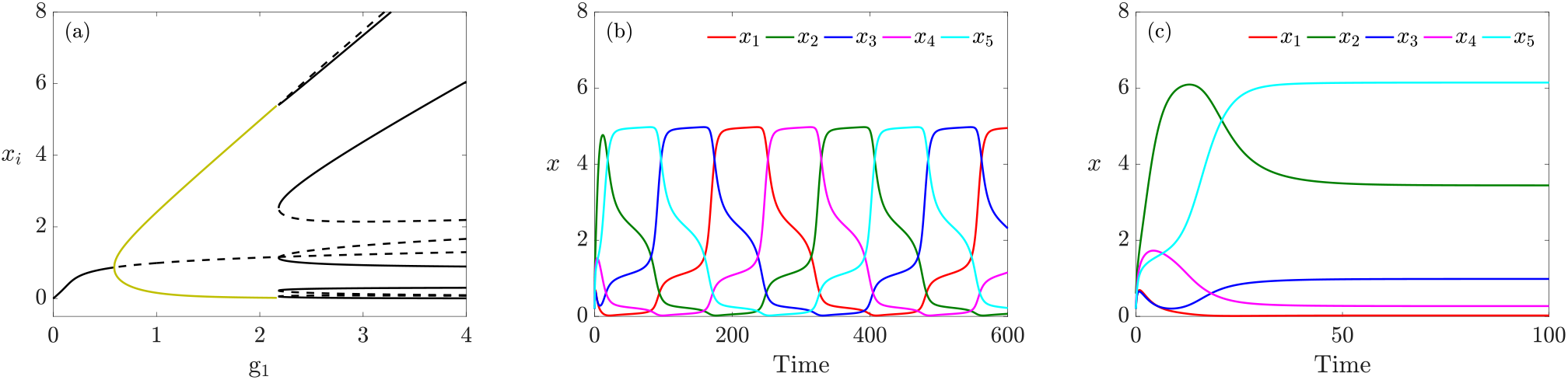
(a) Bifurcation diagram of system (5) with respect to *g*_1_ when *g*_2_ = 0.35, *g*_3_ = 0.2 and *d* = 0.4. (b) Time series of the solutions in the oscillatory/multipotent regime when *g*_1_ = 2. (c), and multistable/ differentiated regime for *g*_1_ = 2.5.

Biologically, the stable generation of cells expressing mixtures of key fate specification TFs shown by these four- and five-gene models matches closely the concept of combinatorial TF expression defining similar cell-types. This is most keenly demonstrated in the peripheral and central nervous systems where distinct neuron types express mixtures of TFs that together specify their individual fate [33]. It may be that the fully differentiated phenotype represents the impact of a mix of TFs expressed at different levels, even where we tend to think of a single dominant master regulator, such as Mitf in melanocyte development.

## 6 Conclusions

We have presented a detailed bifurcation analysis of four variants of the standard repressilator model, and discussed their suitability as novel generic models for the process of cellular differentiation. While the original re-pressilator fails to provide convincing dynamical behaviour, we find that the variants proposed here exhibit biologically relevant features. Our modelling moves beyond Waddington’s vision of the epigenetic landscape since such a picture cannot capture highly dynamical internal states of cells which our model places centre stage.

Bifurcation analysis of the systems using [16] reveals that the oscillatory behaviour emerges from a Hopf bifurcation. At the late stages of multipotency, the oscillating cell visits a set of sub-states where the relevant fate-specific transcription factor(s) are highly expressed relative to the other gene(s). The oscillations terminate at a global bifurcation and the cell settles at a state resembling that of a committed cell-type. From a biological perspective, we can think of the individual fate-specific transcription factors (TFs) as defining individual cell-fates. Where they oscillate, the stem cell retains multipotency (it sequentially transits nearby all states), but at the same time it becomes periodically biased to adopt individual fates due, we propose, to changing exposure to fate-specification signals. When such a signal is received at a sufficiently high level, it may force oscillations to cease and a single TF becomes stably expressed at the expense of the others – cell differentiation has begun.

We have focused on two possible scenarios for driving the oscillatory behaviour. In the first scenario, the same pathway is responsible for initiating and terminating the oscillations, by the increase of the chosen parameter. In the second scenario, a first pathway brings the system in the oscillatory region, and a second one intervenes to drive the exit from it. Our results show that the second scenario, however, gives more control over cell fate. Moreover, the timing of controlling signals is essential in our framework, since this combines with ICs to determine the specific TF that remains stably expressed, and thus the differentiated cell-type.

Depending on the type of gates assumed in the network, one or two genes (or more in networks of higher dimensionality) can be co-expressed during both the oscillatory phase and in the selected differentiated cell type. During the oscillations, combinations of genes may be expressed transiently – these might correspond to partially-restricted cell intermediates (expressing a subset of fate-specific TFs); however, we emphasise that in our scenarios these retain oscillations and hence must be thought of as fully multipotent, despite their ‘snapshot appearance’. In the selected differentiated cell-type, the combination and the levels of expressed TFs vary with ICs and timing of the fate-specification factor; thus different cell-types, with specific quantitatively distinguishable combinations of TFs, are generated. Such a model now provides a view of how multipotency might be exhibited in stem cells, in contrast to the series of bipotential intermediates that underpin standard thinking in development. Note that the external signals drive a fully multipotent cell to a specific committed state directly, without further intermediates with restricted potency. Finally, the case with which stable (committed) states consisting of mixed expression of two or more TFs can be generated may well relate to the mechanism producing related cell-types, e.g. neuron types in both the central and peripheral nervous system.

There is a wider context for the results presented here; the identification of dynamical scenarios typical for repressilator-type GRNs, particularly in view of their proposed role in clock circuits underlying global patterning processes in development [34]. Although the developmental context considered there is different, the formulation and behaviour of the mathematical models has some points of similarity with our analysis, such as the occurrence of Hopf and SNIC bifurcations, which are generic bifurcations involving time-periodic oscillations.

A key difference with our work is ordering of the different regimes. Model 1 of [34] shows a progression from oscillations to a single stable equilibrium, and then a saddle-node bifurcation that creates multiple equilibria. Model 2 of [34] has no Hopf bifurcation but instead a SNIC separating oscillatory from multistable regime; there is no single equilibrium. In contrast, our model robustly indicates a progression from a single equilibrium to oscillations via a Hopf bifurcation, with the oscillations then terminating at a global bifurcation where multiple equilibria emerge. We note that the distinction between the SNIC mechanism and the combination of the global and saddle-node bifurcations is likely to be extremely difficult to detect experimentally due to the high level of resolution required in time, and the effects of noise. It is however mathematically important in terms of comparisons between models.

In terms of model construction a key point of difference is that both Model 1 and Model 2 in [34] are constructed by interpolating, rather artificially, between two mechanisms potentially driven by distinct and unrelated morphogens. Our dynamical model is structurally more robust and parsimonious as our work has shown that the transition from equilibrium to oscillations whose frequency drops to zero, to ‘differentiated’ equilibria, is a natural result of a single GRN mechanism. Naturally, spatial patterning is not part of our model since we focus on the dynamics within a single cell, but establishing connections between temporal dynamics and spatial patterning is an intriguing research direction that we will explore in a future paper.

## Acknowledgments

We gratefully acknowledge Laura Hattam and Michael Thomas for helpful preliminary modelling work. RNK gratefully acknowledges support from the Institute for Mathematical Innovation at the University of Bath that helped to advance this project in its early stages. Funding was provided by BBSRC grants BB/S01604X/1 (SF and AR) and BB/S015906/1 (KCS, JHPD, RNK).

## References

[1] Waddington CH. 1940 Organisers and Genes. Cambridge, UK: Cambridge University Press.

[2] Vibert L, Aquino G, Gehring I, Subkhankulova T, Schilling TF, Rocco A, Kelsh RN. 2016 An ongoing role for Wnt signaling in differentiating melanocytes in vivo. Pigment Cell Melanoma Res. 30, 219–232. (doi:10.1111/pcmr.12568).

[3] Dorsky RI, Raible DW, Moon RT. 2000 Direct regulation of nacre, a zebrafish MITF homolog required for pigment cell formation, by the Wnt pathway. Genes & Dev. 14, 158–162. (doi:10.1101/gad.14.2.158).

[4] Lewis JL, Bonner J, Modrell M, Ragland JW, Moon RT, Dorsky RI, Raible DW. 2004 Reiterated Wnt signaling during zebrafish neural crest development. Development 131, 1299–1308. (doi:10.1242/dev.01007).

[5] Ferrell Jr JE. 2012 Bistability, bifurcations, and Waddington’s epigenetic landscape. Curr. Biol. 22, R458. (doi:10.1016/j.cub.2012.03.045).

[6] Huang S, Li F, Zhou JX, Qian H. 2017 Processes on the emergent landscapes of biochemical reaction networks and heterogeneous cell population dynamics: differentiation in living matters. J. R. Soc. Interface 14, 20170097. (doi:10.1098/rsif.2017.0097).

[7] Huang S. 2010 Cell Lineage Determination in State Space: A Systems View Brings Flexibility to Dogmatic Canonical Rules. PloS Biol. 8, e1000380. (doi:10.1371/journal.pbio.1000380).

[8] Wigger M, Kisielewska K, Filimonow K, Plusa B, Maleszewski M, Suwińska A. 2017 Plasticity of the inner cell mass in mouse blastocyst is restricted by the activity of FGF/MAPK pathway. Sci. Rep. 7, 15136 (doi:10.1038/s41598-017-15427-0).

[9] Graf T, Stadtfeld M. 2008 Heterogeneity of embryonic and adult stem cells. Cell Stem Cell 3, 480–483 (doi:10.1016/j.stem.2008.10.007).

[10] Nikaido M. et al., 2021 Zebrafish pigment cells develop directly from highly multipotent progenitors. Submitted.

[11] Kelsh RN, Camargo Sosa K, Farjami S, Makeev V, Dawes JHP, Rocco A. 2021 Cyclical Fate Restriction, a new view of neural crest cell fate specification. Submitted.

[12] Perez-Carrasco R, Barnes CP, Schaerli Y, Isalan M, Briscoe J, Page KM. 2018 Combining a Toggle Switch and a Repressilator within the AC-DC Circuit Generates Distinct Dynamical Behaviors. Cell Syst. 6, 521–530.e3. (doi:10.1016/j.cels.2018.02.008).

[13] Suzuki N, Furusawa C, Kaneko K, 2011 Oscillatory Protein Expression Dynamics Endows Stem Cells with Robust Differentiation Potential. Plos One 6, e27232. (doi:10.1371/journal.pone.0027232).

[14] Rabajante JF, Babierra AL. 2015 Branching and oscillations in the epigenetic landscape of cell-fate determination. Prog. Biophys. Mol. Biol. 117, 240–249. (doi:10.1016/j.pbiomolbio.2015.01.006).

[15] Kaity B, Sarkar R, Chakrabarti B, Mitra MK. 2018 Reprogramming, oscillations and transdifferentiation in epigenetic landscapes. Sci. Rep. 8, 7358 (doi:10.1038/s41598-018-25556-9).

[16] Doedel EJ, Champneys AR, Fairgrieve T, Kuznetsov YA, Oldeman BE, Paffenroth R, Sandstede B, Wang X, Zhang C. 2010 AUTO-07p: Continuation and Bifurcation Software for Ordinary Differential Equations. Concordia University, Canada. Available from:http://cmvl.cs.concordia.ca/auto.

[17] Elowitz MB, Leibler S. 2000 A synthetic oscillatory network of transcriptional regulators. Nature 403, 335–338. (doi:10.1038/35002125).

[18] Saint-Jeannet JP, He X, Varmus HE, Dawid IB. 1997 Regulation of dorsal fate in the neuraxis by Wnt-1 and Wnt-3a. PNAS 94, 13713–13718. (doi:10.1073/pnas.94.25.13713).

[19] Dorsky RI, Moon RT, Raible DW. 1998 Control of neural crest cell fate by the Wnt signalling pathway. Nature 396, 370–373. (doi:10.1038/24620).

[20] Dunn KJ, Williams BO, Li Y, Pavan WJ. 2000 Neural crest-directed gene transfer demonstrates Wnt1 role in melanocyte expansion and differentiation during mouse development. PNAS 97, 10050–10055. (doi.org/10.1073/pnas.97.18.10050).

[21] Takada R, Takada S, Takeda K, Yasumoto K, Watanabe K, Udono T, Saito H, Takahashi K, Shibahara S. 2000 Induction of Melanocyte-specific Microphthalmia-associated Transcription Factor by Wnt-3a. The Journal of Biological Chemistry 275, 14013–14016. (doi:10.1074/jbc.C000113200).

[22] Jin EJ, Erickson CA, Takada S, Burrus LW. 2001 Wnt and BMP signaling govern lineage segregation of melanocytes in the avian embryo. Dev. Biol. 233, 22–37. (doi:10.1006/dbio.2001.0222).

[23] Lee HY, Kléber M, Hari L, Brault V, Suter U, Taketo MM, Kemler R, Sommer L. 2004 Instructive Role of Wnt/*β*-Catenin in Sensory Fate Specification in Neural Crest Stem Cells. Science 303, 1020–1023. (doi:10.1126/science.1091611).

[24] Greenhill ER, Rocco A, Vibert L, Nikaido M, Kelsh RN. 2011 An Iterative Genetic and Dynamical Modelling Approach Identifies Novel Features of the Gene Regulatory Network Underlying Melanocyte Development. PLoS Genet. 7, e1002265. (doi:10.1371/journal.pgen.1002265).

[25] Coullet P, Gilli JM, Monticelli N, Vandenberghe N. 2005 A damped pendulum forced with a constant torque. Am. J. Phys., 73, 1122–1128. (doi:10.1119/1.2074027).

[26] Andronov AA, Vitt AA, Khaikin SE. 1996 Theory of Oscillators. New York, US: Pergamon.

[27] Tsai TL, Dawes JHP. 2013 Dynamics near a periodically-perturbed robust heteroclinic cycle. Physica D 262, 14–34. (doi:10.1016/j.physd.2013.07.009).

[28] Gambaudo JM. 1985 Perturbation of a Hopf bifurcation by an external time-periodic forcing. J. Diff. Eq. 57, 172–199. (doi:10.1016/0022-0396(85)90076-2).

[29] Marques F, Meseguer A, Lopez JM, Pacheco JR, Lopez JM. 2013 Bifurcations with imperfect SO(2) symmetry and pinning of rotating waves. Proc. R. Soc. A 469: 20120348. (doi:10.1098/rspa.2012.0348).

[30] Strogatz SH. 1994 Nonlinear Dynamics and Chaos. Reading, US: Addison-Wesley.

[31] Thomas R. 1980 On the Relation Between the Logical Structure of Systems and Their Ability to Generate Multiple Steady States or Sustained Oscillations in Numerical Methods in the Study of Critical Phenomena. 180–193. Berlin, Germany: Springer (doi:10.1007/978-3-642-81703-824).

[32] Dawes JHP et al. Dn-symmetric dynamics under weak symmetry breaking. In preparation.

[33] Lloret-Fernández C, Maicas M, Mora-Martínez C, Artacho A, Jimeno-Martín Á, Chirivella L, Weinberg P, Flames N. 2018 A transcription factor collective defines the HSN serotonergic neuron regulatory landscape. eLife 7, e32785. (doi: 10.7554/eLife.32785).

[34] Jutras-Dubé L, El-Sherif E, François P. 2020 Geometric models for robust encoding of dynamical information into embryonic patterns. eLife 9, e55778. (doi:10.7554/eLife.55778).

