## Supplementary material for "Novel Generic Models for Differentiating Stem Cells Reveal Oscillatory Mechanisms": FCDKR_Suppl_Revised.pdf

Saeed Farjami, Karen Camargo Sosa,
Jonathan H.P. Dawes, Robert N. Kelsh, Andrea Rocco

#### S1 Model Derivation

Following [1], we here provide a derivation of the Hill functions dynamics utilised in the construction of the models in this paper.

##### S1.1 Regulation by repressive transcription factors

Many processes are involved in the regulation of a protein. Here, we consider only few of them and assume that the other processes such as transcription and mRNA translation are implicit. The model includes (un)binding of the transcription factor (TF) $R$, degradation of the protein $P$, and synthesis of $P$ when gene $G$ is in the active state, denoted $G^*$, as follows.

$$R + e_R \underset{\alpha_1}{\overset{\alpha_0}{\rightleftharpoons}} r_R, \qquad G^* \xrightarrow{g} P, \qquad P \xrightarrow{d_P} \emptyset, \tag{S1}$$

where $e_R$ and $r_R$ represent the 'empty' and 'regulated' regulatory elements of $G$, respectively. The parameters over/under the arrows indicate the rate of the processes.

If $[\cdot]$ denote the number of a chemical per cell, the processes in (S1) are described with

$$\begin{cases} \dfrac{d[r_R]}{dt} = \alpha_0[R](1-[r_R]) - \alpha_1[r_R], \\ \dfrac{d[P]}{dt} = g[e_R] - d_P[P]. \end{cases} \tag{S2}$$

where $[e_R] + [r_R] = 1$ and $[G^*] = [e_R]$.

Binding and unbinding of $R$ to the regulatory element is significantly faster than the protein regulation and degradation ($\alpha_0$, $\alpha_1 \gg g$, $d_P$). By quasi-steady-state approximation $d[r_R]/dt = 0$, we have

$$\frac{d[P]}{dt} = g\frac{1}{1+\alpha[R]} - d_P[P], \tag{S3}$$

where $\alpha = \alpha_o/\alpha_1$.

Here, the TF $R$ regulates the protein $P$ as a monomer giving rise to a Hill coefficient one. Hill functions with higher coefficients can be derived when regulation proceeds through binding/unbinding of complexes (dimers, trimers, etc).

##### S1.2 Regulation by two repressors arranged in an 'OR gate'

Now, we consider regulation of gene $G$ by two independent and repressive TFs, $A$ and $B$. In an OR gate configuration, the presence of one TF is sufficient for the repression of $G$. Similar to the previous section, we have

$$A + e_A \underset{\alpha_1}{\overset{\alpha_0}{\rightleftharpoons}} r_A \quad \text{and} \quad B + e_B \underset{\beta_1}{\overset{\beta_0}{\rightleftharpoons}} r_B \tag{S4}$$

with steady states

$$[e_A] = \frac{\alpha_1}{\alpha_1 + \alpha_0[A]} = \frac{1}{1+\alpha[A]} \quad \text{and} \quad [e_B] = \frac{\beta_1}{\beta_1 + \beta_0[B]} = \frac{1}{1+\beta[B]}. \tag{S5}$$

where $\alpha = \alpha_0/\alpha_1$ and $\beta = \beta_0/\beta_1$.

Because $A$ and $B$ are repressing and act independently, gene $G$ remains active when both binding sites are empty. Accordingly,

$$[G^*] = [e_A][e_B] = (1 - r_A)(1 - r_B). \tag{S6}$$

Therefore, the dynamics of the protein $P$ is described by

$$\frac{d[P]}{dt} = g\frac{1}{1+\alpha[A]} \cdot \frac{1}{1+\beta[B]} - d_P[P]. \tag{S7}$$

##### S1.3 Regulation by two repressors arranged in an 'AND gate'

In an AND gate configuration, the presence of both TFs $A$ and $B$ is necessary for the repression of $G$. Accordingly, gene $G$ is active when at least one of the binding sites is empty. In this case, the average number of activated genes is

$$[G^*] = [G_1^*] + [G_2^*] + [G_3^*] = [e_A][e_B] + [r_A][e_B] + [e_A][r_B]. \tag{S8}$$

where $[r_A]$ and $[r_B]$ are given by equation (S5). For generality, we here assume that the production rate of protein $P$ depends on the configuration of bound TF [2]. Therefore, by assuming production rates $g_1$, $g_2$ and $g_3$ for $[G_1^*]$, $[G_2^*]$ and $[G_3^*]$, respectively, the dynamics of protein $P$ is described by

$$\frac{d[P]}{dt} = g_1\frac{1}{1+\alpha[A]} \cdot \frac{1}{1+\beta[B]} + g_2\frac{\alpha[A]}{1+\alpha[A]} \cdot \frac{1}{1+\beta[B]} + g_3\frac{1}{1+\alpha[A]} \cdot \frac{\beta[B]}{1+\beta[B]} - d_P[P]. \tag{S9}$$

This can be rewritten as

$$\frac{d[P]}{dt} = \frac{g_1 + g_2\alpha[A] + g_3\beta[B]}{(1+\alpha[A])(1+\beta[B])} - d_P[P], \tag{S10}$$

which is the form used in (3) and (5).

##### S1.4 Regulation by three repressors arranged in an 'AND gate'

Similarly to the AND gate with two repressors, derived above, the average number of activated genes for the AND gate with three repressors is

$$[G^*] = [e_A][e_B][e_C] + [e_A][e_B][r_C] + [e_A][r_B][e_C] + [r_A][e_B][e_C] + [e_A][r_B][r_C] + [r_A][e_B][r_C] + [r_A][r_B][e_C]. \tag{S11}$$

Accordingly, assuming different maximal expression rate for each configuration of binding gives

$$\frac{d[P]}{dt} = \frac{g_1 + g_2\alpha[A] + g_3\beta[B] + g_4\gamma[C] + g_5\beta\gamma[B][C] + g_6\alpha\gamma[A][C] + g_7\alpha\beta[A][B]}{(1+\alpha[A])(1+\beta[B])(1+\gamma[C])} - d_P[P], \tag{S12}$$

which is the form used in (4).

#### S2 Analysis of the oscillation period in the standard repressilator

Here we show a numerical exploration of the dependency of the period of oscillations of the standard repressilator on the two significant parameters in the model. We restrict attention to the fully-symmetric version for simplicity: $g_1 = g_2 = g_3$, $d_1 = d_2 = d_3$, $\alpha_1 = \alpha_2 = \alpha_3$ and $b_1 = b_2 = b_3$ in all cases. We consider separately the variation in the period $T$ of the oscillation with changes in $g_1$ and in $d_1$; these parameters are as defined in (1). Our simulations are presented in Fig. S1.

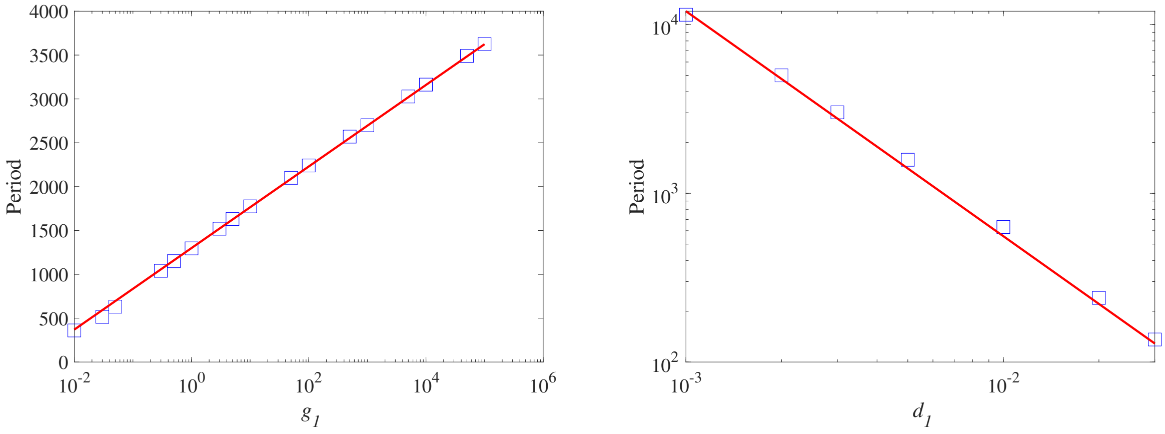


Figure S1: **Fit of the oscillations period dependency on $g_1$ and $d_1$ parameters in the repressilator model.** In both panels the points (blue squares) represent the result of the simulations for $d_1 = d_2 = d_3 = 0.01$ fixed (left) and $g_1 = g_2 = g_3 = 0.05$ fixed (right), while the solid lines correspond to best fitting curves as detailed in the text. The resulting fitting parameters are $a_0 = 1300$, $a_1 = 202$, $a_2 = 1.2$, and $\beta = 4/3$. In all cases $b_1 = b_2 = b_3 = 10^{-5}$ and $\alpha_1 = \alpha_2 = \alpha_3 = 50$.

The fitting functions assumed in these cases are as follows:

$$T(g_1) = a_0 + a_1 \log(g_1) \qquad \text{for } d_1 \text{ fixed} \tag{S13}$$

and

$$T(d_1) = a_2 d_1^{-\beta} \qquad \text{for } g_1 \text{ fixed} \tag{S14}$$

The simulations show a very weak dependency of the period on the parameter $g_1$ - over seven orders of magnitude in $g_1$ the period increases by a factor of only around seven. As $d_1$ increases the period drops much more rapidly, in line with the power-law scaling $T \propto d_1^{-4/3}$. This is to be expected since the degradation rate $d_1$ is a key timescale in the problem and the decay rate of the concentrations of TFs is essential to the nature of the 'repressilator' dynamics. In neither case does the qualitative nature of the periodic oscillation change over these ranges of parameters. This allows us to conclude that the parameter values used, for example in figure 2 in the main text, are entirely typical of the repressilator's dynamical behaviour. Finally we note that the dependence of the period and shape of the oscillation on the production rates $b_1, b_2$ and $b_3$ is very weak in this parameter range where typical concentrations of TFs are maintained well above each of the $b_i$ by the nonlinear prodution terms.

### S3 Analysis of mathematical concepts in the cross-repressilator

Three mathematical concepts that need to be explained and analysed more closely are symmetric structure of systems (2) and (3), the growth rate of period of limit cycles close to SNIC/heteroclinic bifurcation and determining factors in the order of expressed genes when the system oscillates.

#### S3.1 Analysis of the three-gene Models in the symmetric case

In the case that the models (2) and (3) are cyclically symmetric under the permutation $(x_1, x_2, x_3) \to (x_2, x_3, x_1)$, i.e. when the parameters $g$, $\alpha$, $\beta$, $k$ and $h$ do not take different values in each equation, the symmetric structure of the problem enables some semi-analytic progress to be made since the Jacobian matrix at the symmetric equilibrium point $(x, x, x)$ is constrained to always take the form

$$Df|_{(x,x,x)} = \begin{pmatrix} a & b & c \\ c & a & b \\ b & c & a \end{pmatrix}, \tag{S15}$$

where the matrix entries are the partial derivatives of the function on the right-hand side, say $\dot{x}_1 = f_1(x_1, x_2, x_3)$. In particular we have

$$a \equiv \left.\frac{\partial f_1}{\partial x_1}\right|_{(x,x,x)}, \qquad b \equiv \left.\frac{\partial f_1}{\partial x_2}\right|_{(x,x,x)}, \quad \text{and} \quad c \equiv \left.\frac{\partial f_1}{\partial x_3}\right|_{(x,x,x)}, \tag{S16}$$

where the partial derivatives are evaluated at the symmetric equilibrium point which can usually not be solved for explicitly. The eigenvalues of Section S3.1 are then a single real eigenvalue and a complex conjugate pair

$$\lambda_1 = a + b + c, \tag{S17}$$

$$\lambda_{2,3} = a - \frac{1}{2}(b+c) \pm \frac{i\sqrt{3}}{2}(b-c). \tag{S18}$$

In the case of the original repressilator model (1) for which $f_1(x_1, x_2, x_3) = b + \frac{g}{1+\alpha x_2^h} - kx_1$, we see that the location of the equilibrium point satisfies $\frac{g}{1+\alpha x^h} = kx - b$. Its eigenvalues are therefore

$$\lambda_1 = -k - \frac{\alpha h}{g} x^{h-1}(kx-b)^2, \tag{S19}$$

$$\lambda_{2,3} = -k + \frac{\alpha h}{2g} x^{h-1}(kx-b)^2 \pm \frac{i\sqrt{3}\alpha h}{2g} x^{h-1}(kx-b)^2. \tag{S20}$$

So $\lambda_1 < 0$ always, and at the Hopf bifurcation point, where $Re(\lambda_{2,3}) = 0$ we see that the frequency $Im(\lambda_{2,3}) = k\sqrt{3}$ independent of the other parameters. In the symmetric case this implies that the frequency at the two Hopf bifurcations from the symmetric equilibrium, illustrated in Fig. 4 (left), would be equal.

For the cross-repressilator with the 'OR' gate we have $f_1(x_1, x_2, x_3) = \frac{g}{(1+\alpha x_2^h)(1+\beta x_3^h)} - kx_1$. Hence the symmetric equilibrium $(x, x, x)$ satisfies $(1+\alpha x^h)(1+\beta x^h) = \frac{g}{kx}$. Computing the relevant partial derivatives yields the three eigenvalues

$$\lambda_1 = -k - \frac{k\alpha h x^h}{1+\alpha x^h} - \frac{k\beta h x^h}{1+\beta x^h} \tag{S21}$$

$$\lambda_{2,3} = -k + \frac{1}{2}\left(\frac{k\alpha h x^h}{1+\alpha x^h} + \frac{k\beta h x^h}{1+\beta x^h}\right) \pm \frac{i\sqrt{3}k^2 h(\beta-\alpha)x^{1+h}}{2g} \tag{S22}$$

which shows that in this case also, $\lambda_1 < 0$ and there is a possibility of a Hopf bifurcation where $Re(\lambda_{2,3}) = 0$. The imaginary part of the complex conjugate pair is proportional to $(\alpha - \beta)$ which indicates, in agreement with numerical explorations, that these eigenvalues become real along the line $\alpha = \beta$.

Finally we note that the occurrence of the SNIC bifurcation and the codimension-two point indicated by the grey dot in Fig. 6 are also consequences of symmetry. We do not comment further here, and leave a more detailed analysis of the consequences of symmetry for this problem to be the subject of a future paper [3].

#### S3.2 Analysis of the oscillation period close to a global (SNIC/heteroclinic) bifurcation

In this subsection we investigate in more detail the scaling behaviour of the increase of the period of the limit cycles as they terminate at two different global bifurcations: the saddle-node on an invariant circle (SNIC) bifurcation and the heteroclinic bifurcation. Appropriate plots of the period of the orbit near these global bifurcations yields straight-line plots. For both of these bifurcations, the period increases in accordance with a well-known analytical scaling law when $g \longrightarrow g_{\text{SNIC}}$. Close to a SNIC bifurcation, it is known that $T \propto 1/\sqrt{|g - g_{\text{SNIC}}|}$, i.e. a straight line in log-log scale. The left panel in Fig. S2 shows a transformation of Fig. 4(b) in log-log scale which therefore matches with what we expect for a SNIC bifurcation. For $\alpha = 9$ we find $g_{\text{SNIC}} \approx 1.51449$.

When the periodic orbit terminates at a heteroclinic bifurcation, the period increases according to $T \propto \log|g - g_{\text{HC}}|$ as $g \longrightarrow g_{\text{HC}}$. Therefore a plot of the period $T$ against $g - g_{\text{HC}}$ on a log-linear scale should be a straight line. We confirm this in the right panel of Fig. S2 where we show the data computed for Fig. 6 (right) but on a log-linear scale, again shifted by subtraction of $g_{\text{HC}} \approx 1.05509$.

A further detailed verification of our numerical work is that the theoretical result for the constant in the relation $T \propto \log 1/|g - g_{\text{HC}}|$ is given by $3/\lambda_+$, i.e. $T \sim (3/\lambda_+)\log(1/|g - g_{\text{HC}}|)$ where $\lambda_+$ is the positive eigenvalue of

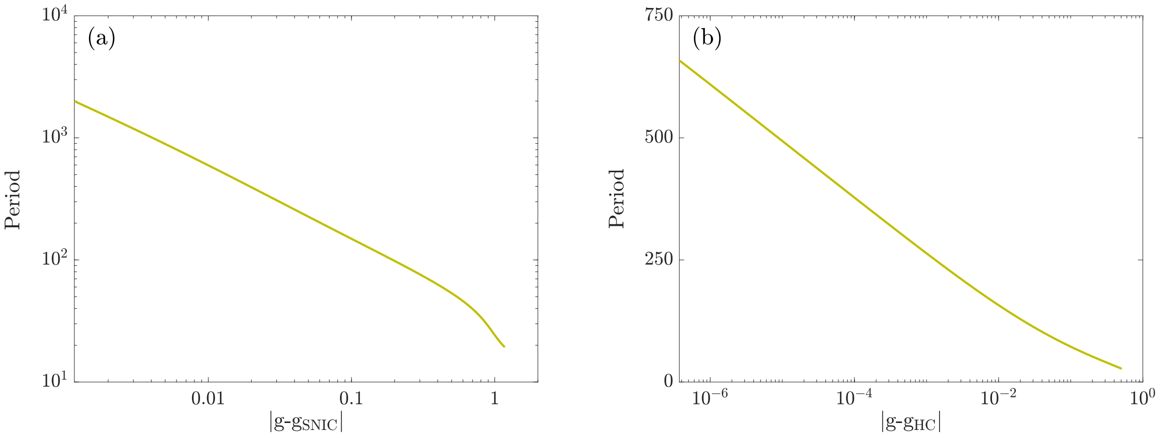


Figure S2: **Growth scale of periodic orbits.** Period of the family of periodic orbits ending at a SNIC bifurcation for $\alpha = 9$ (left) and heteroclinic bifurcation for $\alpha = 1$ (right). The rest of parameters of the system are the same as Fig. 4(b).

the saddle point equilibrium that collides with the periodic orbit (in the case that the periodic orbit is stable, which it is here). The factor of 3 arises due to the $\mathbb{Z}_3$ symmetry and the three distinct saddle points that all collide with the periodic orbit simultaneously. Numerically, we find that $\lambda_+ \approx 0.05922$ (5 dp); therefore $1/\lambda_+ = 16.88619$ (5 dp). Hence the theoretical estimate for the slope of the straight line is $3 \times 16.88619 = 50.65857$ (5 dp). From the right panel of Fig. S2 we estimate the slope of the curve to be 50.17112 based on the values of the period T at $|g - g_{HC}| = 3 \times 10^{-7}$ and $|g - g_{HC}| \approx 3 \times 10^{-4}$. This therefore provides quantitative confirmation of the bifurcation behaviour in the differential equations.

##### S3.3 Ordering of the gene expression in the oscillations

As discussed in the main text, the genes are expressed in a particular order in the oscillatory regimes. For instance, in the three-gene case with the OR gate configuration, the gene expression occurs cyclically clockwise. That is, specifically, first gene $X_3$ is expressed and then genes $X_2$ and finally $X_1$. This order of gene expression is due to the values of the parameters of (2). In (2), we chose $\alpha = 9 > \beta = 0.1$ which makes the inhibitions of the inner loop in Fig. 3 stronger; this results in the genes being expressed in clockwise order. However, if we set $\alpha < \beta$, the order of gene expression in the oscillatory regime is reversed.

Mathematically the oscillatory direction of the family of periodic orbits emanating from a (supercritical) Hopf bifurcation is determined by the relative signs of the real and imaginary parts of the eigenvectors corresponding to the pair of complex conjugate eigenvalues ($\lambda_{2,3} = Re(\lambda_{2,3}) \pm Im(\lambda_{2,3})$). Since these eigenvectors remain non-zero and vary continuously along the Hopf bifurcation curve on which these eigenvalues have zero real part, we can conclude that the oscillation changes direction when the imaginary parts of the eigenvalues pass through zero. Figure S3 shows the imaginary parts of $\lambda_{2,3}$ and its conjugate $-\lambda_{2,3}$ along the curve of Hopf bifurcations (dark green) in Fig. 6 projected on $g$ and $\alpha$ (in log scale) axes. For small values of $g$ (large values of $\alpha$), let assume that $Im(\lambda_{2,3})$ is positive (blue); therefore, its conjugate, $-Im(\lambda_{2,3})$, is negative (red). As $g$ increases ($\alpha$ decreases), $Im(\lambda_{2,3})$ and its conjugate, $-Im(\lambda_{2,3})$, cross each other. Thus, the direction of oscillation of periodic orbits switches. The crossing point of $Im(\lambda_{2,3})$ and $-Im(\lambda_{2,3})$ at 0 corresponds to the point that the Hopf curve (HB) touches the homoclinic curve (HC, red). Precisely at the parameter value where the complex conjugate pair of eigenvalues are zero, the Hopf bifurcation theorem breaks down and further analysis is required; this discussion will be presented in future work [3].

#### S4 Numerical Methods

We have employed different tools and software packages for computing solutions, bifurcation diagrams and colourmaps. All the time series have been integrated in XPPAUT, freely available at `http://www.math.pitt.`

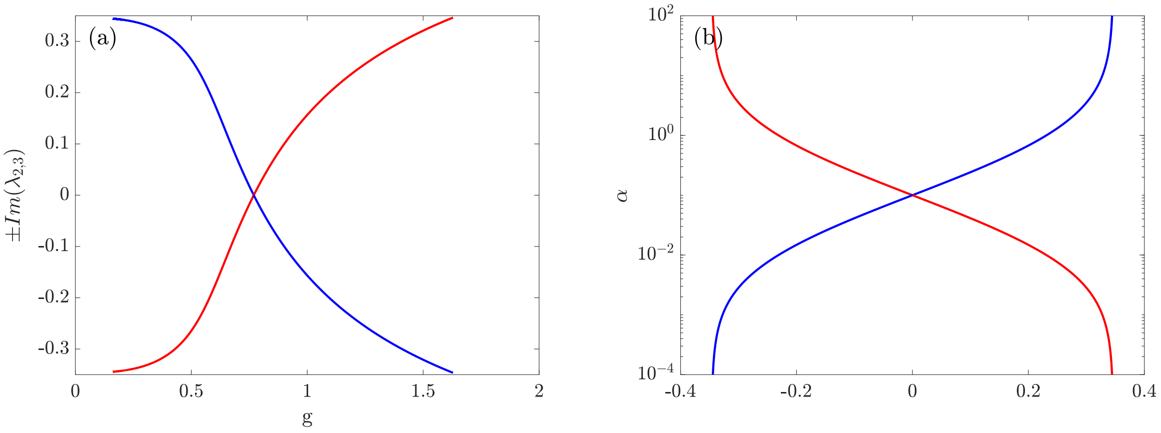


Figure S3: **Imaginary parts of eigenvalues along the Hopf curve.** The variation in the imaginary parts of the complex conjugate eigenvalues along the Hopf bifurcation curve (HB, dark green) in Fig. 6 shown versus $g$ (left panel) and $\alpha$ in log scale (right panel). Other parameter values are as in Fig. 6.

edu/~bard/xpp/xpp.html. The bifurcation diagrams are computed using pseudo-arclength continuation methods in software package AUTO [4,5].

Finally, the colourmaps are computed in MATLAB® (Mathworks Inc.) starting from different initial conditions. In Fig. 5, the initial-condition planes are colour-coded depending on the steady state (one of the three) that the system converges to. Similarly, in Fig. 8, the final state of the system when $g$ increases and saturates at a value larger than $g_{\text{SNIC}}$ determines the colour of the initial conditions.

### References

[1] Greenhill ER, Rocco A, Vibert L, Nikaido M, Kelsh RN. 2011 An Iterative Genetic and Dynamical Modelling Approach Identifies Novel Features of the Gene Regulatory Network Underlying Melanocyte Development. PLoS Genet. **7**, e1002265. (doi:10.1371/journal.pgen.1002265).

[2] Andrecut M, Halley JD, Winkler DA, Huang S. 2011 A General Model for Binary Cell Fate Decision Gene Circuits with Degeneracy: Indeterminacy and Switch Behavior in the Absence of Cooperativity. PLoS ONE **6**, e19358. (doi:10.1371/journal.pone.0019358).

[3] Dawes JHP, Farjami S, et al, $D_n$-symmetric dynamics under weak symmetry breaking. In preparation.

[4] Doedel EJ. 1981 AUTO: A Program For the Automatic Bifurcation Analysis of Autonomous Systems. Congr. Numer. **30**, 265–284.

[5] Doedel EJ, Oldeman BE. AUTO-07p: Continuation and Bifurcation Software for Ordinary Differential Equations. Department of Computer Science, Concordia University,Montréal, Canada; 2010. With major contributions from A.R. Champneys, F. Dercole, T.F. Fairgrieve, Y. Kuznetsov, R.C. Paffenroth, B. Sandstede, X.J. Wang and C.H. Zhang. Available from: http://www.cmvl.cs.concordia.ca.
